# Cerebellar neurodynamics during motor planning predict decision timing and outcome on single-trial level

**DOI:** 10.1101/833889

**Authors:** Qian Lin, Magdalena Helmreich, Friederike Schlumm, Jennifer M. Li, Drew N. Robson, Florian Engert, Alexander Schier, Tobias Nöbauer, Alipasha Vaziri

**Author notes:** Corresponding author: Alipasha Vaziri, Laboratory of Neurotechnology and Biophysics, The Rockefeller University, 1230 York Avenue, New York, NY 10065.

## Abstract

The neuronal basis of goal-directed behavior requires interaction of multiple separated brain regions. How subcortical regions and their interactions with brain-wide activity are involved in action selection is less understood. We have investigated this question by developing an assay based on whole-brain volumetric calcium imaging using light-field microscopy combined with an operant-conditioning task in larval zebrafish. We find global and recurring dynamics of brain states to exhibit pre-motor bifurcations towards mutually exclusive decision outcomes which arises from a spatially distributed network. Within this network the cerebellum shows a particularly strong pre-motor activity, predictive of both the timing and outcome of behavior up to ∼10 seconds before movement initiation. Furthermore, on the single-trial level, decision directions can be inferred from the difference neuroactivity between the ipsilateral and contralateral hemispheres, while the decision time can be quantitatively predicted by the rate of bi-hemispheric population ramping activity. Our results point towards a cognitive role of the cerebellum and its importance in motor planning.

## INTRODUCTION

Animals interact with their environment in a goal-directed manner and through complex behaviors that are optimized over time. Such goal-directed behaviors require motor planning that precedes actions. The neuronal signatures of such preparatory activity have been reported in the cortex, in particular in the motor cortex (Churchland et al., 2010), pre-motor cortex (Li et al., 2016) and parietal cortex (Maimon and Assad, 2006). They display a few common features (Gold and Shadlen, 2007; Murakami and Mainen, 2015; Shenoy et al., 2013; Svoboda and Li, 2018): First, preparatory activity can occur long before movement initiation. Examples include humans engaged in a self-initiated binary decision tasks (Soon et al., 2008), mice during an instructed-delay task (Li et al., 2015), or primates involved in a visuomotor task (Maimon and Assad, 2006), where preparatory activity encoding for decision outcomes has several seconds preceding movement initiation has been found in various cortical areas. Secondly, preparatory neuroactivity has been shown to manifest itself as a ramping or sequential activity to a threshold (Harvey et al., 2012) which can be found across different brain regions, contexts and animal models and the rate of which has been shown to be correlated with the decision timing (Gold and Shadlen, 2007; Maimon and Assad, 2006; Mazurek et al., 2003; Murakami and Mainen, 2015; Murakami et al., 2014; Wang, 2008; 2012). Finally, the neuroactivity during the pre-motor period can on the level of individual neurons take more complex forms than merely a sub-threshold version of movement-related activity (Churchland and Shenoy, 2007). For instance, in a delayed-reach task, the tuning of individual neurons to preferred direction in the motor and pre-motor cortex has been shown to differ during pre-motor and movement epochs of the task (Churchland et al., 2010).

These observations are incompatible with static and trial-averaged tuning frameworks for representation of motor responses, as averaging can mask the inherent trial-to-trial variability of neuronal activity and motor response, and thus their relationship on a trial-by-trial basis (Afshar et al., 2011; Briggman et al., 2005; Churchland and Kiani, 2016; Wei et al., 2019). Frameworks based on dynamical systems—in which the movement-related neuronal dynamics are described by differential equations whose initial conditions are set by the preparatory activity—have been shown to provide a better model than trial-averaged representation frameworks for how motor behavior emerges from pre-motor neuronal population dynamics (Churchland et al., 2010; 2012; Mante et al., 2013; Michaels et al., 2016; Pandarinath et al., 2018; Shenoy et al., 2013).

While the above findings in the cortex have provided some insights into the neuronal basis underlying action selection and the relationship between preparatory neuroactivity and motor response, several observations point towards the importance of taking a brain-wide approach to decision making that also includes subcortical regions and their interactions with the cortex (Cisek and Kalaska, 2010; Gao et al., 2018; Guo et al., 2017; Horwitz and Newsome, 1999; Kunimatsu et al., 2018; Liu et al., 2013). Support for this notion comes also from the fact that decision making almost inevitably requires access to or can be modulated by brain functions such as short-term memory, internal states or reward expectation that are known to have a sub-cortical representation (Abbott et al., 2017; Allen et al., 2017). However, obtaining large-scale, high spatio-temporal resolution access to neuroactivity of cortical as well as sub-cortical regions is currently not technically possible in mammalian brains. On the other hand, studying complex goal-directed behaviors, motor planning and decision making in invertebrate model systems is typically difficult and findings may not be directly translatable to the mammalian brain.

Larval zebrafish represents an attractive model to investigate the above questions due to its high level of physiological and genetic homology to the mammalian brain, conformity to basic vertebrate brain organization (Friedrich et al., 2010; Kalueff et al., 2014) and its transparent brain that allows for whole-brain *in vivo* optical recording of neuroactivity at high speed and cellular resolution (Ahrens et al., 2012; 2013; Cong et al., 2017; Favre-Bulle et al., 2018; Migault et al., 2018; Nöbauer et al., 2017; Portugues et al., 2014; Prevedel et al., 2014; Wolf et al., 2015). However, establishing a robust behavioral paradigm in the larval zebrafish that involves learning, memory and integration of information across different brain regions in the context of more cognitive task and within which the neuronal basis of goal directed motor planning and decision making could be studied, has proven challenging.

In the present study, we have combined high-speed volumetric whole-brain calcium imaging based on light-field microscopy (LFM) (Nöbauer et al., 2017; Prevedel et al., 2014) with a new learning paradigm in larval zebrafish for operant conditioning, Relief Of Aversive Stimulus by Turn (ROAST) (Li, 2013), to investigate the neuronal basis of action selection and the underlying pre-motor activity. In LFM (Broxton et al., 2013; Cong et al., 2017; Levoy et al., 2006; Nöbauer et al., 2017; Pégard et al., 2016; Prevedel et al., 2014), different positions within a volume in the sample are mapped via a microlens array onto different patterns on the camera sensor plane which are all acquired in parallel. Acquisition is followed by computational reconstruction to obtain the three-dimensional images of the sample. Other calcium imaging techniques used in larval zebrafish work by sequentially scanning a diffraction limited point (Stosiek et al., 2003) or a light-sheet (Ahrens et al., 2013) across the sample volume, which limits their volume rates. In contrast to these techniques, LFM provides a volumetric imaging rate that allows for simultaneous and volumetric recording of the activities of neurons on the whole-brain level. As a result, LFM opens up the possibility to study a qualitatively different range of questions on the single-trial level, as discussed above (Churchland and Shenoy, 2007) and in which circuit-level detail can be analyzed across entire neuronal systems. The present study represents to our knowledge the first application of LFM to a neuroscientific question in vertebrates.

In ROAST (Li, 2013), fish learn to perform a directed tail movement to the right or the left in order to receive relief from an aversive heat stimulus. While the ROAST assay has been initially developed to study learning (Li, 2013), it also provides an ideal opportunity to investigate the neuronal basis of action selection and the underlying pre-motor activity. The binary nature of the decision as well as the extended decision time, between the heat onset and the turning behavior, allow studying the neuronal dynamics that underlie the choice for the turn direction as well as the timing for movement initiation. Using LFM while animals were engaged in ROAST provided us with a brain-wide model for the representation and interaction of different brain regions that are involved in representation of sensory information, learning, reward prediction, as well as the generation of decision-related pre- and post-motor neuroactivity.

We found brain states (i.e., the state of activity of all neurons at a given time), to display stimuli-triggered recurrences as well as a decision-related bifurcation in brain-state space at the trial-by-trial level. While the heat stimulus was a main driver of global brain state transitions, decision outcomes could be also decoded from the global brain dynamics, suggesting involvement of multiple brain areas. We found preparatory neuroactivity preceding the motor response in multiple parts of the fish brain. Among these, the cerebellum exhibited a surprisingly strong level of preparatory activity as well as the adjacent anterior rhombencephalic turning region (ARTR), a self-oscillating hindbrain population, also known as hindbrain oscillator (HBO), which has been shown to introduce a bias in the turn direction during exploratory locomotion (Dunn et al., 2016; Wolf et al., 2017). The activity from these regions allowed us to predict the turn direction about 10 seconds before turn initiations. The presence of preparatory activity in the cerebellum points towards the critical role that it could be playing in motor planning more generally, and, as zebrafish lack a cortex, more specifically prior to emergence of cortical regions and functions (Hodos, 2009). Furthermore, we found that the neural population dynamics of the cerebellum and ARTR were able to explain two key aspects of our decision task at the single-trial level: the turn direction and its timing. The outcome of the turn direction was determined by an ipsi-contra-lateral competition of neuroactivity, which was more prominent in animals that learned the task than non-learner animals, while the timing of the turn could be quantitatively predicted at single-trial level by the collective ipsi-contra lateral ramping rate of neuronal population dynamics in the cerebellum and the ARTR. Thus, our work extends the traditional view on the role of the cerebellum in motor control and provides support for its more cognitive functions. While our observations are in line with previously proposed “ramping-to-threshold” models (Gold and Shadlen, 2007; Wang, 2012), our results extend this framework to single-trials eliminating the need of trial-averaging which has been previously necessary to detect neuronal ramping activities (Shadlen et al., 2016). Further, our data-driven model provides the basis for a framework for decision making based on competition-cooperation in which both, the competition - i.e. the differential neuroactivity - and corporation - i.e. the joint neuroactivity - of two ipsi- and contralaterally located neuronal populations can simultaneously encode different aspects of the decision.

## RESULTS

### An operant conditioning paradigm of larval zebrafish

We aimed to find a robust behavioral paradigm that involved learning and short-term memory while exhibiting a delay period from the onset of an instructing sensory cue to the execution of motor response, during which the neuronal basis of motor planning and decision making could be studied. While previously used assays in larval zebrafish for sensorimotor transformation lack the above features, most of the typically used assays in rodents or primates study motor planning by introducing a delay period from the onset of the stimulus to a go cue, after which the animal is trained to initiate a motor response (Mohebi and Oweiss, 2014; Shenoy et al., 2013; Svoboda and Li, 2018). However, motor responses of animals engaged in naturalistic action selections are self-initiated and happen in the in the absence of a go cue. The ROAST operant conditioning paradigm (Figure 1A) addresses both issues. In each trial of this task, head-fixed larval zebrafish are first exposed to a mildly aversive heat stimulus generated by an infrared laser source that is directed at their head region (see STAR Methods). Each fish is exposed to two training blocks with opposite reward directions in which animals have to perform a turn by moving their tail towards a specific direction, i.e. left or right, in order to obtain relief from the heat stimulus (Figure 1A). Data of an example learner fish that learned in both training blocks is shown in Figure 1B. After a few incorrect turns at the beginning of each block, the animal learned the correct reward direction (see STAR Methods for details).

**Figure 1.**
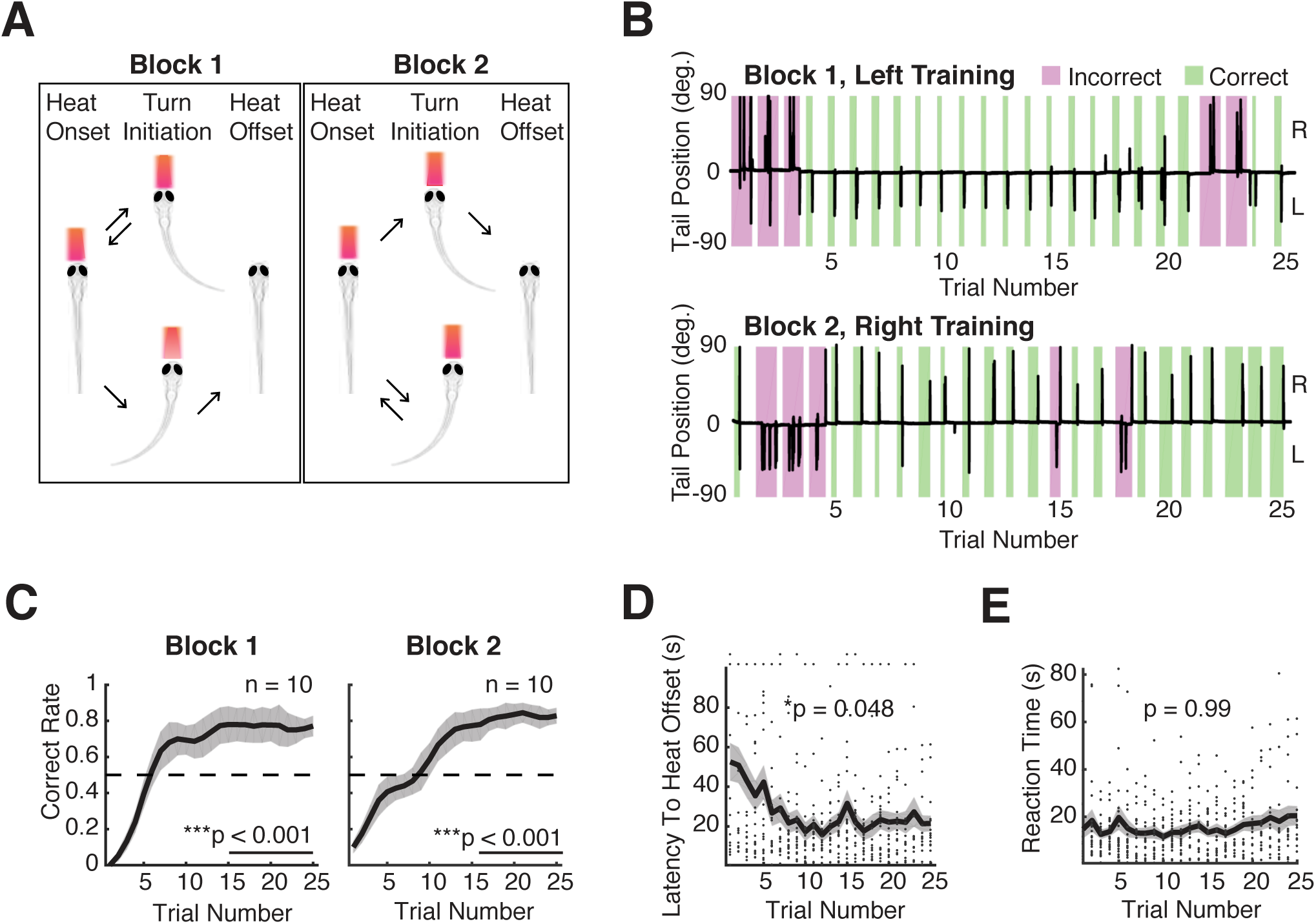
An operant conditioning assay for larval zebrafish: Relief of Aversive Stimulus by Turn (ROAST). (A) Relief of Aversive Stimulus by Turn (ROAST) in head-fixed larval zebrafish. Head-fixed, tail free larval zebrafish receive a mildly aversive heat stimulus by an infrared laser (red trapezoid) at the beginning of each trial. The laser is turned off if the fish makes a tail movement in the reward direction and remains on otherwise. In the second training block the reward direction is switched, with each block consisting of 20-25 trials (see STAR Methods for details). (B) The learning progress (two blocks) of an example learner fish (8 dpf). Black traces indicate tail positions as a function of time, magenta and green rectangles indicate the duration of the heat stimulus for each incorrect and correct trial. (C) Averaged performance as a function of trial number from 7-8 dpf larvae (n = 10) (mean ± SEM). The correct rate in the first 10 trials is significantly below 0.5, **p = 0.001 for Block 1 and *p = 0.012 for Block 2; the correct rate in the last 10 trials is significantly above 0.5, ***p < 0.001 for both blocks (p based on bootstrapping, n = 5000). (D) Decrease of exposure to heat as a function of trials, averaged from both blocks of 10 learners (mean ± SEM). Dots indicate single trials. Two tailed *p = 0.048, Kruskal-Wallis test. Black dots indicate individual trials. (E) Reaction time, defined as the time from heat onset to the first movement, as a function of trial number remains stable, averaged from both blocks of 10 learners (mean ± SEM). Dots indicate single trials. Two-tailed p = 0.99, Kruskal-Wallis test.

We performed simultaneous whole-brain calcium imaging using a home-built LFM (Nöbauer et al., 2017; Prevedel et al., 2014) and a transgenic fish line *Tg(elavl3:H2B-GCaMP6s)* (Dunn et al., 2016) with pan-neuronal expression of nucleus-localized GCaMP6s (NL-GCaMP6s). The inherent mapping of sample volume to 2D-image sensor of our LFM allowed us to capture the activity states of all neurons on the whole-brain level within a volume of (700 μm × 700 μm × 200 μm), all voxels acquired truly simultaneously and at a volume rate of 10 Hz. The learners’ performance became stable typically after the first 15 trials in both blocks, with an asymptotic final performance of 80% correct (Figure 1C), while the non-learners showed overall correct rates well below 50% (Figure S1A and see STAR Methods). At the level of single trials, we measured the latency to heat offset, the time from onset of the heat stimulus until the first correct turn, and the reaction time (RT), the time between the onset of heat stimulus and the initiation of the first movement, regardless of it being correct or incorrect. The latter is widely used to quantify the decision time (Wang, 2012). We found that the latency to heat offset was reduced as a function of trials in learners (Figure 1D), but not in non-learners (Figure S1B). In contrast, we found a relatively stable mean RT, independent of the training progress in both learners and non-learners (Figure 1E, S1C). These results demonstrate that larval zebrafish can learn to perform a directed tail movement to obtain relief from an aversive heat stimulus. This capability of the animals, together with the design of the ROAST assay—which provides a sufficiently long delay period between the stimulus onset and the movement initiation—opens up the possibility to study the neuronal basis of preparatory motor activity in the context of a binary decision task on the whole-brain level.

### Recurring brain state dynamics capture representation of heat and show bifurcation for different behavioral outcomes

Volumetric reconstruction of the LFM data (see also Movie S1), followed by neuronal signal extraction using our established data processing pipeline (Prevedel et al., 2014), allowed us to obtain neuronal activity traces from ∼5000 of the most active neurons across the entire brain. Figure 2 shows the data for an example learner (Block 2, right training). When these activities were co-registered with the time period of heat stimulus and when the time points of directed left or right tail movement, a significant level of correlation was observed (Figure 2A). Mapping these neuronal activities spatially onto the fish brain revealed active neurons across the entire forebrain, midbrain and the anterior hindbrain. We found distinct neuronal dynamics associated with different brain regions that were correlated with specific epochs of our task (Figure 2B). In correct trials, a “Heat ON” event triggered neuronal activity in the telencephalon (white and red arrow heads, Figure 2B(i)), followed by an increase in activity of the ipsilateral cerebellum and a decrease in thalamic activity (yellow arrow heads and white arrow, Figure 2B (ii)). At turn initiation, the ipsilateral cerebellum (white arrow, Figure 2B (iii)) and ARTR, i.e. HBO, (white arrow head) were highly active, while after heat offset, the activity of the ipsilateral cerebellum (white arrow, Figure 2B (iv)), but not that of ARTR (white arrow head, Figure 2B (iv)), decreased and at the same time the contralateral cerebellum and ARTR increased their activity (Figure 2B (iv)). In parallel also an increase of activity in the thalamus and the telencephalon was observed in the period between 1-15 second after a “Heat OFF” event (yellow arrow heads, Figure 2B (iv)). This increased activity level decayed over longer time periods after “Heat OFF” (Figure 2B (v)).

**Figure 2.**
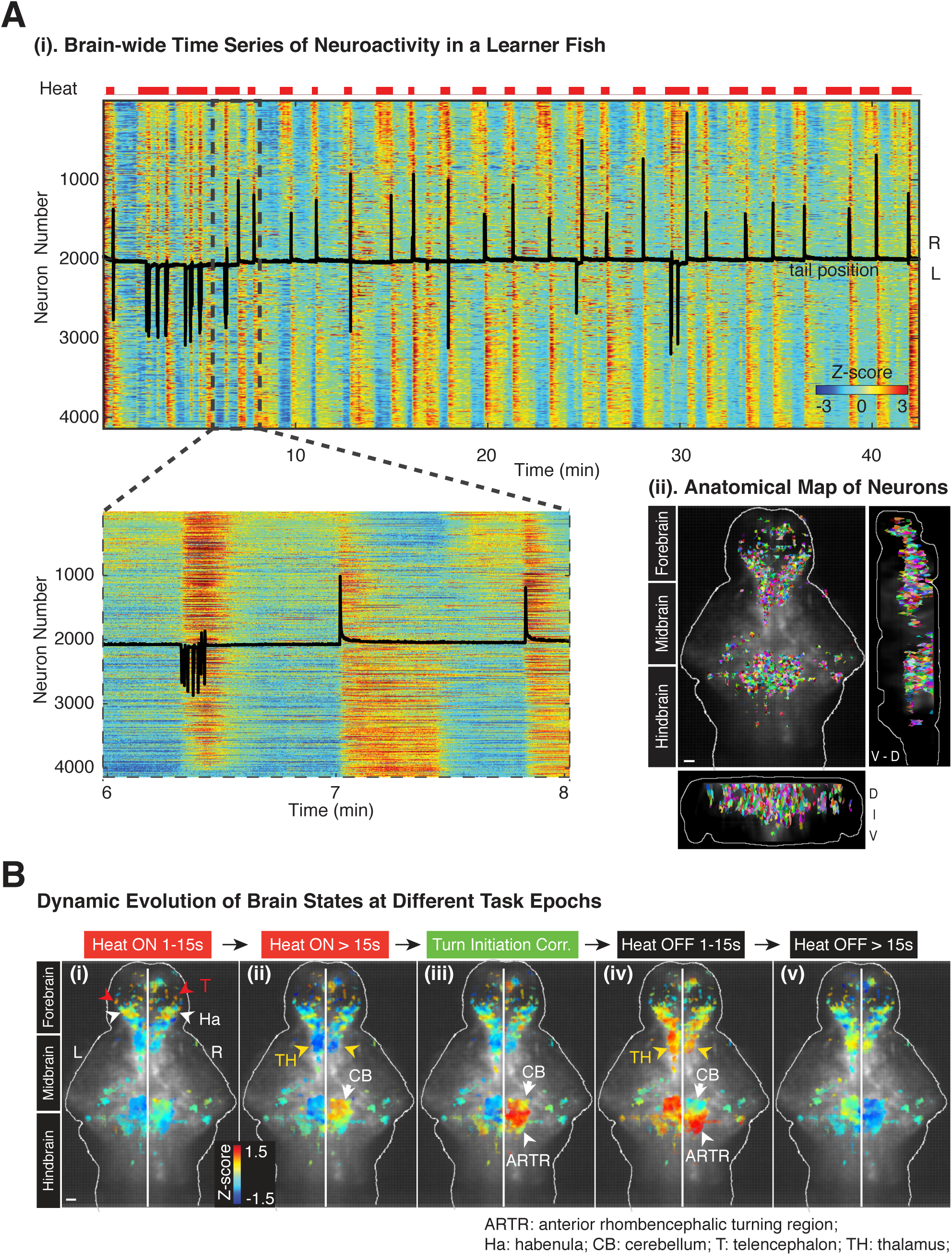
Whole-brain dynamics of neuronal activity and behavior during different task epochs. (A) (i) Heat map of brain-wide neuronal time series (Z-scored) from a learner with top 2% active neurons extracted, resulting in 4124 neurons. Neurons sorted by their correlations with heat stimuli, from highly positive to highly negative. Upper red bars: heat stimuli; superimposed black trace: tail position of the fish; lower panel: zoom in of the dashed rectangular area. (ii) Anatomical map of these exacted neurons. Neurons are with different colors. Scale bar, 50 µm. (B) Dynamic evolution of brain states, i.e. the activity of all neurons at a given time point, at different task epochs. (i) During the initial 15 s of Heat ON, the forebrain neurons show increased activity; white arrow heads: habenula (Ha); red arrow heads: telencephalon (T). (ii) For Heat ON > 15 s, the activity in the ipsilateral cerebellum (CB, white arrow) increases, while the thalamus (TH, yellow arrow heads) decreases. (iii) At correct turn initiation, the ipsilateral cerebellum and ARTR are highly active (white arrow and arrow head respectively). (iv) During initial 15 s of Heat OFF, the activity of the ipsilateral cerebellum (white arrow), but not ARTR (white arrow head), decreases, while that of the contralateral cerebellum, ARTR and the thalamus (yellow arrow heads) increases. (v) All neuroactivity reaches the baseline levels again for longer periods after Heat OFF. Color-coding shows averaged brain states during same epochs across trials; spatial filters of neurons are maximum-projected and superimposed on average fluorescence for anatomical reference. Same dataset as panel (A). Scale bar, 50 µm.

We next set out to identify how the dynamics of brain states — i.e., the state of activity of all ∼5000 neurons across the entire brain at a given time— would be associated with the different task epochs by performing a linear dimensionality reduction based on principal component analysis (PCA). However, PCA on brain states failed to effectively reduce dimensionality in a fashion that would capture key differences in dynamics during different epochs of our paradigm (Figure S2A) and would facilitate a simple visualization in two or three dimension as has been done in the past for smaller nervous systems, such as that of *C. elegans* (Kato et al., 2015). This was also consistent with the observation that the first three PCs captured <60% of the variance of the brain dynamics (Figure S2B).

To overcome this issue, we applied a nonlinear dimensionality reduction technique t-distributed stochastic neighbor embedding (t-SNE) to the time series of the brain states (Maaten and Hinton, 2008). Briefly, t-SNE transforms the structure contained in high-dimensional data into a two- or three-dimensional map by minimizing the Kullback-Leiber divergence between the Gaussian similarity matrix of the data points and the t-distributed similarity matrix of the map points. t-SNE maintains the local structure by keeping similar data points close to each other on the t-SNE map (Maaten and Hinton, 2008). Even though t-SNE has the disadvantage of not providing the same visualization on each run of the optimizer, nor an explicit transformation matrix that would be required to establish a quantitative correspondence between the t-SNE map and neuroanatomy, it nevertheless provides an effective, exploratory tool for identifying hidden, low-dimensional structure in high-dimensional data (Davie et al., 2018; Grün and van Oudenaarden, 2015; LeCun et al., 2015; Macosko et al., 2015).

Applying t-SNE to our brain states allowed us to identify, on the highest level, two major clusters of brain states that were associated with the presence and absence of the heat stimulus respectively (Figure 3A). Consistent with this observation, quantitative analysis of the t-SNE map revealed that the inter-cluster distance, i.e. Euclidian distance of embedded brain states between the “Heat ON” and the “Heat OFF” cluster was on average significantly higher than the intra-cluster distance of brain states within each cluster for our example fish (Figure 3B) as well as across fish on the population level (Figure S3A). These results show that changes of the heat stimulus are the driver of the temporally sharp transitions between brain states, which underlie the most prominent feature of the t-SNE map.

**Figure 3.**
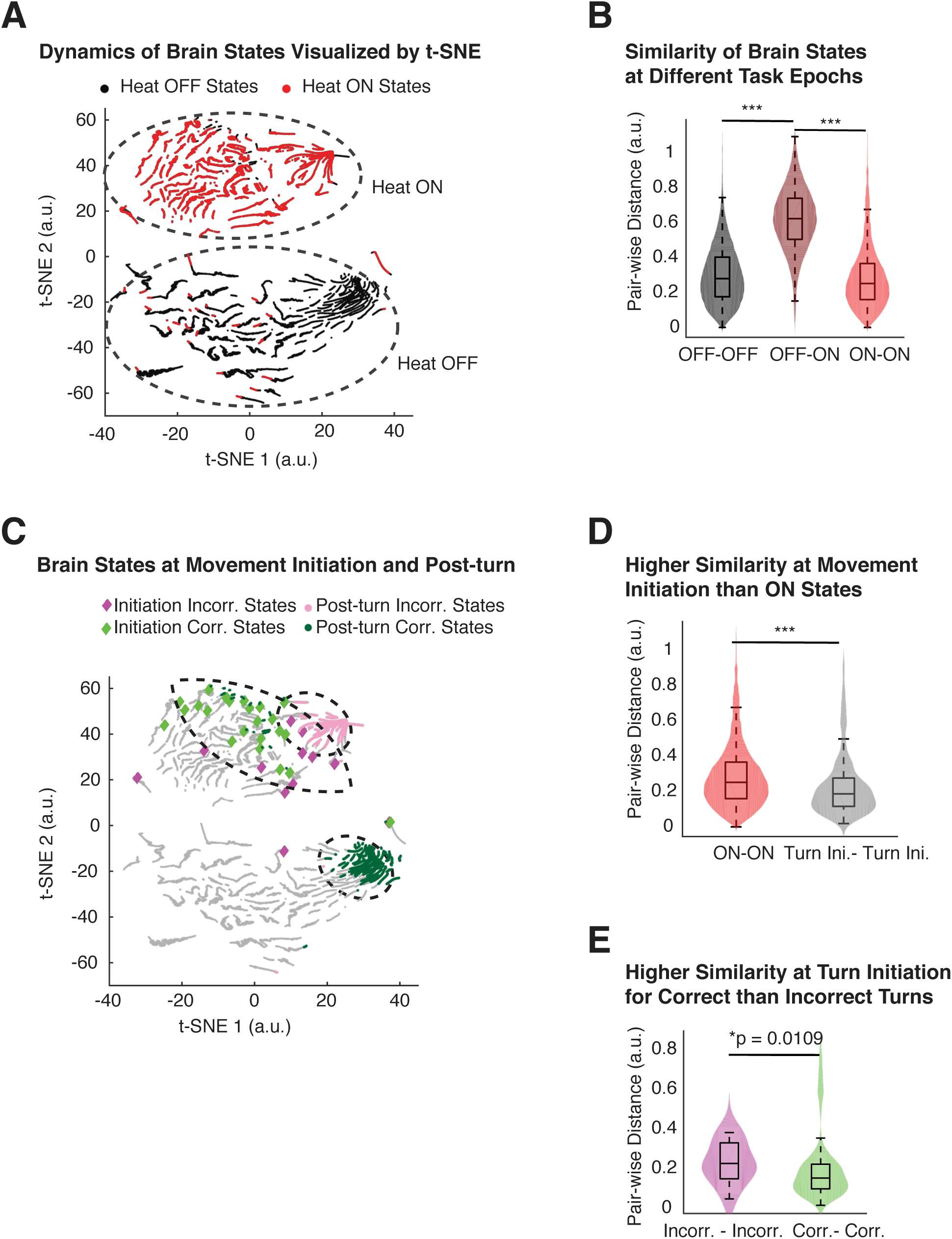
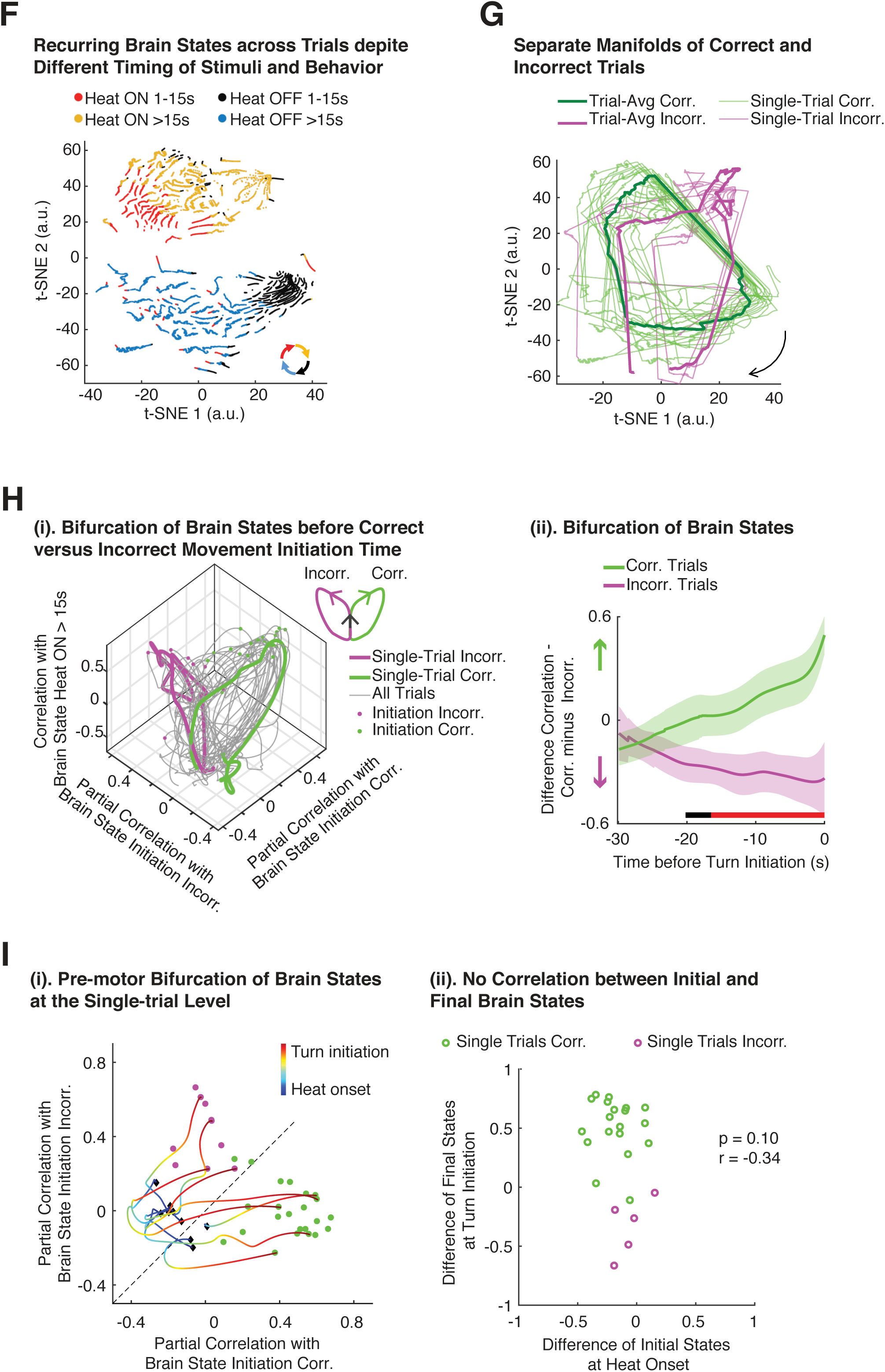
Dynamics of the brain states show recurrence to stimuli and bifurcation at different behavioral outcomes. (A) Dynamic evolution of brain states visualized in a 2D t-SNE map, with major transitions in brain states associated with Heat ON and Heat OFF. Brain states (n = 24472) represented as dots, color-coded by the stimuli. (B) Similarity of brain states within and across Heat ON and OFF. Within-epoch brain states show a higher level of similarity than across-epoch, calculated as the pair-wise distances between brain states; example data ***p < 0.001, Wilcoxon rank sum test. Same for multiple animals (n = 12, Figure S3A). (C) Convergence of brain states at turn initiation and post turn (0-10 s) evolution; color-code: Incorrect (Incorr.): magenta, correct (Corr.): green. Turn initiations occur during Heat ON, and the corresponding brain states are located close to each other within a sub-region of the “Heat ON” cluster, with few exceptions. Post-incorrect turns remain within the “Heat ON” cluster while post-correct turns transition to the “Heat OFF” cluster. (D) Brain states at movement initiation exhibit a higher level of similarity than brain states on average with the “Heat ON” cluster, ***p < 0.001, Wilcoxon rank sum test. Same for multiple animals (n = 12, Figure S3B). (E) Brain states resulting in correct turns exhibit a higher level of similarity at movement initiation than those resulting in incorrect turns, Wilcoxon rank sum test. Same for multiple animals (n = 12, Figure S3C). (F) Brain states reoccur across trials, forming a loop despite differences in timing of stimulus and motor response. Color-coded by the different task epochs. (G) Temporal evolution of brain states for the correct and incorrected trials results in separate cyclic manifolds. Hausdorff distance (Bruno et al., 2017), ***p < 0.0001, Wilcoxon rank sum test. Thick lines, averages and thin lines, single trials; arrow indicates direction of temporal evolution. (H) Bifurcation of brain states before turn initiation. (i) After heat onset, all brain states exhibit similarity along “Heat ON” dimension (vertical axis), followed by a pre-turn bifurcation towards the correct or incorrect state. Similarity is measured by partial correlation between a given brain state and the average correct, incorrect or “Heat ON” state (see Methods for details). (ii) Bifurcation of brain states occurs 20 s before turn initiation. Brain states is measured as the difference in correlation with the average value for correct brain states and with the average value for incorrect brain states. Brain states are aligned to the time point of movement initiation. Solid lines and shaded bands represent mean ± 95% CI. Wilcoxon rank sum test, two-tailed. Time points with *p < 0.05 marked with horizontal black bar, those with **p < 0.01 with red bar. Similar bifurcation of brain states before turn initiation can be observed in other fish (n = 4). (I) 2D representation of single-trial bifurcation process. (i) Individual correct and incorrect trials are highlighted during the pre-motor period (Black diamonds: heat onset, green and magenta dots: turn initiations for correct and incorrect turns, respectively). See also Movie S2. (ii) No correlation between brain states at heat onset and the brain states at turn initiation, Spearman’s correlation. Brain states characterized using the same metric as in H(ii).

Next, we investigated the representation and the changes of brain states that were induced by decision-related behaviors. As expected, we found that post-correct-turn brain states were embedded within the “Heat OFF” cluster while the post-incorrect-turn brain states were part of the “Heat ON” cluster (Figure 3C). Moreover, they each densely and distinctively occupied a sub-region of each cluster (Figure 3C), indicating the high level of similarity between brain states across trials immediately after a correct and incorrect motor response, respectively. Further, before a turn event, we found that the similarity between brain states across trials was significantly higher than the average similarity between brain states in the “Heat ON” cluster (Figure 3D, S3B). This pre-turn similarity of brain states was significantly higher for trials that resulted in a correct turn than those leading to an incorrect turn (Figure 3E, S3C). These observations show that the brain repeatedly assumes global states that are consistent across trials; that sequences of these global states are to a certain degree stereotypical (both after and before motor behavior); and that immediately before movement initiation these global brain states exhibit a lower variability in trials that result in a correct behavior.

To explore the above point further we investigated how sequences of brain states might encode information on performed or planned motor behavior. To do so, we color-coded the brain states in both, the “Heat ON” and the “Heat OFF” cluster according to task epochs. In both clusters, the brain states evolved smoothly as a function of time and in a cyclic manner (Figure 3F). This was better visualized on the single-trial level by connecting temporally consecutive brain states in the t-SNE map (Figure 3G). We found the brain state dynamic to form a cyclic manifold, with the correct and incorrect trials forming distinct loops (Figure 3G). Using the Hausdorff metric to quantify the pairwise distance between the single-trial loops, we found the pairwise distance between pairs of correct trials was significantly smaller than the distance between pairs of incorrect trials or between correct-incorrect trial pairs (***p < 0.0001, Wilcoxon rank sum test). This shows that during correct trials, highly similar brain states occur across trials, despite the different duration of the stimulus in each trial and the different timing of the decisions.

To better visualize the temporal evolution of brain states in each trial towards the decision time point, and to highlight the differing evolution in correct versus incorrect trials, we calculated the time-dependent correlation (Pearson’s r) between each brain state in a trial and a reference brain state that was calculated by averaging the brain states of the same type (i.e., correct versus incorrect), as they displayed a high level of similarity across trials (Figure 3C). Moreover, because brain states at movement initiation were part of “Heat ON” clusters, we used partial correlation coefficients (see STAR Methods) to factor out the heat-induced neuronal contribution both in correct and incorrect initiation states (Figure 3H). The brain state first displayed increases similarity along the “Heat ON” dimension, and then shows a bifurcation towards the correct or incorrect decision states (Figure 3H (i)) which on the single-trial level occurred about 20 seconds before turn initiations (*p < 0.05, Figure 3H (ii)). Such single-trial bifurcation towards binary decision direction are shown for different reaction times in Figure 3I (i) and Movie S2. We found no correlation (Spearman’s rho = −0.34, p = 0.10) between the initial brain states at the onset of the heat stimulus and the movement initiation brain states (Figure 3I (ii)).

### Signatures of pre-motor bifurcating neuroactivity distributed across the brain but highly concentrated in cerebellum and ARTR

Given our observations on the level of brain states, we set out to investigate the brain regions that gave rise to the activities of the neurons constituting the brain states that drove the bifurcation towards different decision outcomes. Such information cannot be directly extracted from our t-SNE analysis of the brain states as t-SNE learns only a non-parametric mapping (Maaten and Hinton, 2008). Thus, we applied t-SNE onto neuronal time series in order to partition them first into functional clusters and then searched for decision-dependent signals in each cluster.

To do so we embedded regressors of the behavior and the heat stimulus into the neuronal space as “baits’”. We defined 4 regressors: “TurnL”, “TurnR”, “Heat ON” and “Heat OFF”, and obtained them by convolving the actual time series of each regressor with the temporal response kernel of our calcium sensor (NL-GCaMP6s) (see STAR Methods). Figure 4 shows data for a right training in the second block of a learner fish, with corresponding regressors shown in Figure 4A. Applying t-SNE on the time series (∼30,000 time points) of all neurons (∼5,000) and that of our five regressors allowed us to identify the individual functional neurons and their clustering around our regressors, which we emphasized further by partitioning the t-SNE map using hierarchical clustering (see STAR Methods and Figure 4B).

**Figure 4.**
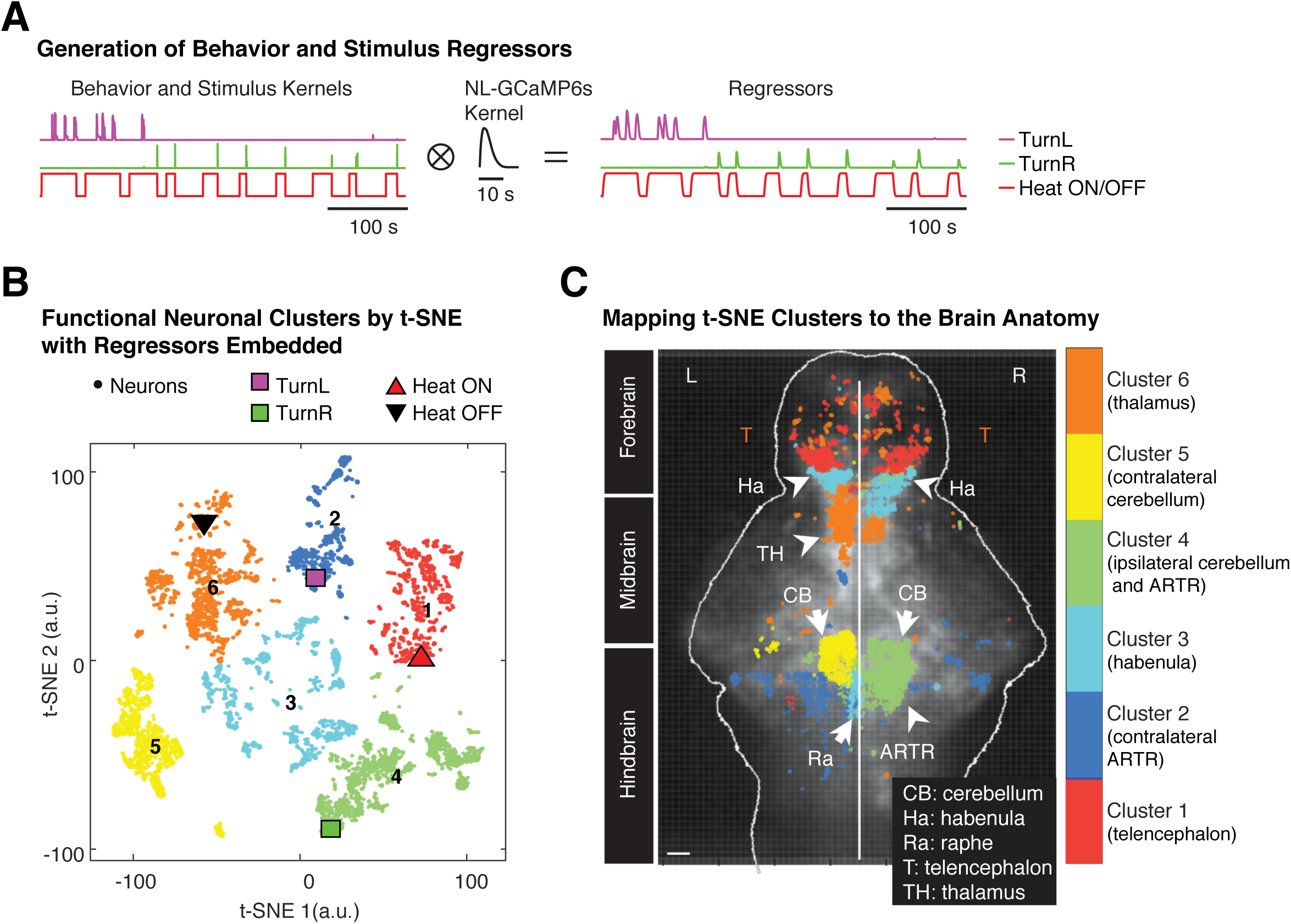
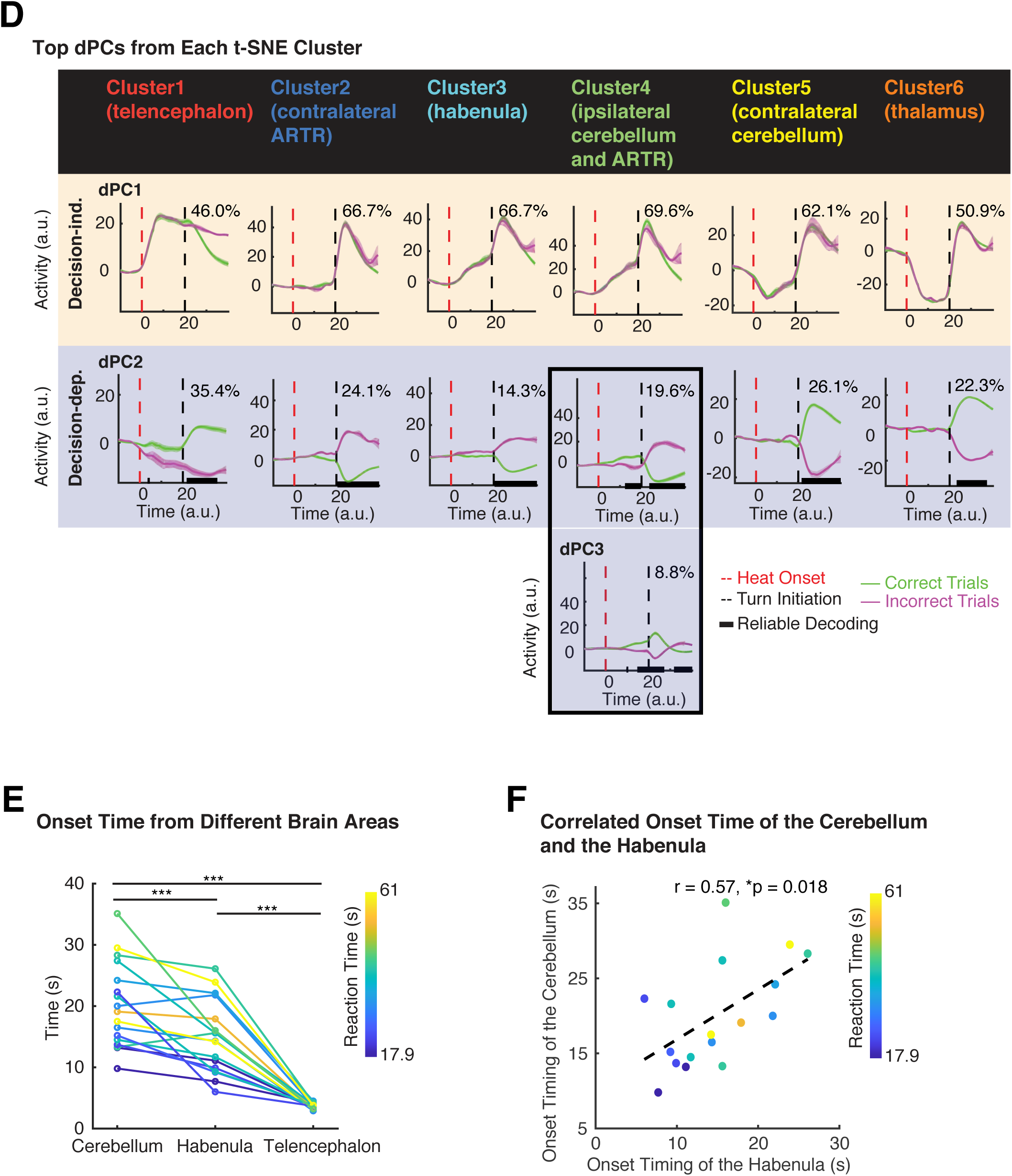
Pre-motor bifurcation activity is spatially-distributed across the entire brain, yet highly concentrated in the cerebellum & ARTR neurons. (A) Example behavior and stimulus regressors obtained by convolving behavior and stimulus time traces with the nucleus-localized (NL)-GCaMP6s kernel. These regressors were fed into t-SNE together with neuronal activity in order to obtain meaningful interpretations for neuron clusters obtained via t-SNE. (B) Distinct functional clusters of neurons with the behavior and stimulus regressors embedded in a 2D t-SNE map. This t-SNE map is partitioned by hierarchical clustering. The Heat ON regressor is located in Cluster 1, Heat OFF in Cluster 6, TurnR in Cluster 4, TurnL in Cluster 2. No regressors in Cluster 3 and Cluster 5. (C) Anatomical distribution of these t-SNE clusters. Cluster 1 is mainly in the telencephalon (T). Cluster 2 in the left side of hindbrain, overlapping with the left ARTR, consistent with the embedding of TurnL regressor. Cluster 3, located in the center of the t-SNE map, consists of the habenula (Ha) and the raphe (Ra). Cluster 4, where the TurnR regressor is embedded, consists of the right cerebellum (CB) and ARTR. Cluster 5 is the left cerebellum (CB). Cluster 6, containing the Heat OFF regressor, is mainly in the thalamus (TH) and sparsely in the telencephalon (T). Scale bar, 50 µm. (D) Top demixed principal components (dPCs) from each cluster. Population activity from each cluster was projected onto individual dPCs and averaged over trials (lines and shaded bands: mean ± 95% CI). Explained variance for each dPC given at top right of each panel. The top two dPCs contain >80% variance of each cluster, except for Cluster 6 with 73% (see Figure S5A). Horizontal black bars mark time points when correct versus incorrect trials can be reliably decoded. Pre-motor decision-dependent signals (in which decoding time precedes and lasts until turn initiation time) from the cerebellum and ARTR highlighted with black box. All clusters show post-turn decoding within second dPC, but only Cluster 4 shows pre-motor decoding, with ∼20% explained variance. Third dPC of Cluster 4 also shows pre-motor decoding for decision directions, while third dPCs in other clusters are decision-independent (not shown). Time series processed for co-alignment (see STAR Methods). (E) Comparison of neuroactivity onset time after stimulus, showing only participating brain areas. The observable differences in onset times imply direction of sensory information flow from telencephalon via habenula to cerebellum. (F) Correlation between neuroactivity onset time in cerebellum and habenula, Pearson’s r; data points are individual trials. Same data as panel (E).

Using this approach, six different well-separated neuronal clusters with five regressors embedded within them were found (Figure 4B). We then marked the anatomical location of each neuron with a color indicating the cluster it belongs to, and found that the t-SNE clusters also formed localized clusters in the fish brain that coincided well with distinct anatomical regions, including the telencephalon, habenula, thalamus, cerebellum and raphe (Figure 4C).

We found a distinct group of bilaterally located neurons (Cluster 1) in the telencephalon that responded to the onset of the heat stimulus, “Heat ON”. Another distinct group (Cluster 6), overlapping largely with the thalamus, but also with neurons sparsely distributed in the telencephalon, encoded for “Heat OFF” (Figure 4 B, C). “TurnL” was represented by contralaterally located neurons in the hindbrain and the left ARTR (Cluster 2), while “TurnR” was encoded by neurons in the ipsilateral right hindbrain that highly overlapped with the cerebellum and the ARTR and fell into a single cluster (Cluster 4). Cluster 3 overlapped with habenula and raphe, while Cluster 5 coincided with the left cerebellum. Our results corroborate previous findings, since the anatomical locations of Cluster 2 and Cluster 4 coincide with the left and right ARTRs which have been found to modulate the probability of left versus right turns (Dunn et al., 2016). The functional clusters that overlapped with the cerebellum and ARTR were not symmetrically distributed: The ipsilateral side was captured by one cluster (Cluster 4) while the neuroactivity of the same anatomical region on the contralateral side was distributed among two clusters (Cluster 2 and Cluster 5). This asymmetry which reflects the higher degree of correlated neuroactivity on the ipsilateral side could result from a stronger input into the ARTR from the ipsilateral cerebellum during the pre-motor period and which acts an instructing signal during learning. This notion was consistent with our observations on the lack of such positive unilateral correlation of the activity of ARTR and the cerebellum in animals engaged in spontaneous turns as well as previous reports on exploratory locomotion (Dunn et al., 2016) and phototaxis (Wolf et al., 2017) in which no learning is involved.

We then searched for the neural activity signals within each cluster that contributed to the observed pre-motor bifurcation of brain states and thus would be *predictive* of the motor response. However, since neurons can in principle have a mixed response, i.e. also decision-independent components such as those associated with the representation of the stimulus, it was necessary to parse out the decision-specific component from each cluster. To do so, we used an extended version of PCA, the demixed principal component analysis (dPCA) (Kobak et al., 2016). dPCA decomposes the activity of a neuronal population into a few task-related components, such as different stimuli and decisions (Kobak et al., 2016). Thereby, besides identifying the transformation and components that account for the majority of the observed variance in the data as in PCA, dPCA can also reveal the dependence of the neural activity on task-related parameters such as stimuli states and decisions. Thus, we configured dPCA to decompose the neuronal population activity in each cluster into decision-dependent versus decision-independent dPCs (see the illustration of the workflow in Figure S4A, B). We then searched the top dPCs of each cluster which together explained ∼80% of observed variance (Figure S5A) for decision-dependent signals (Figure S5B) and asked whether they were able to separate correct from incorrect decisions before movement initiations (Figure 4D).

We found the dPC1 of all clusters to account for at least ∼50% of the observed variance in the corresponding cluster, and to exhibit a decision-independent response. Consistent with our observation of the t-SNE map where the “Heat ON” regressor is embedded in Cluster 1 (telencephalon), we found the dPC1 of Cluster 1 to show increased activity only after heat onset. dPC1 of Cluster 2 (contralateral ARTR) showed increased activity only after turn initiation, suggesting it is encoding for movements, while dPC1 of Cluster 3 (habenula) and Cluster 4 (ipsilateral cerebellum and ARTR) displayed a linear increase in activity upon heat onset followed by a further increase and a secondary peak after turn initiation, suggesting their potential involvement in sensorimotor transformation. dPC1 of Cluster 5 (contralateral cerebellum) and Cluster 6 (thalamus) showed an overall decreased activity upon heat onset – in contrast to the neuroactivity in Cluster 4 and Cluster 1 respectively – followed by an increase of activity and a secondary peak after turn initiation. Given the large overlap of Cluster 5 and Cluster 6 with the contralateral cerebellum and the thalamus respectively, this could result from an ipsilateral inhibition of the cerebellum in the case of Cluster 5 and from an inhibitory innervation from the telencephalon in the case of Cluster 6, or alternatively a local interaction with a pool of inhibitory neurons within thalamus (Halassa and Acsády, 2016).

dPC2 explained ∼25% of observed variance in each cluster. Importantly, we found that it exhibited in all clusters a decision-dependent behavior with a clear separation of correct versus incorrect decisions, in some clusters prior to and in some cluster after initiation of motor behavior (Figure 4D and see STAR Methods). Among these clusters, Cluster 4, the ipsilateral cerebellum and ARTR, allowed for a reliable decoding of the turn direction well before turn initiation (∼8 seconds). A similar pre-motor decision-dependent signature, although with a reversed sign of post-turn activity, was also observed in dPC3 of Cluster 4 (Figure 4D). Overall, even though the explained variance of the entire brain by Cluster 4 was comparable to other clusters (Figure S4C), these two dPCs of Cluster 4 accounted for most (78%) of the observed pre-motor decision-dependent activity over the entire brain (Figure S4D). Other brain regions such as the habenula and raphe (Cluster 3), contralateral cerebellum (Cluster 5), the thalamus (Cluster 6), and the telencephalon (Cluster 1) also exhibited a pre-motor decision-dependent activity in their higher dPCs, i.e. dPC3 and higher dPCs (Figure S4D). In contrast, when we looked at their learning-dependent signals by dividing all trials into pre-and post-learning and then applying dPCA to decompose the activity of each neuron into learning-dependent and learning-independent components, we found that these brain regions, i.e. the thalamus (Cluster 6), the telencephalon (Cluster 1) and the habenula (Cluster 3), accounted for a combined 88% of the variance (Figure S5E), suggesting that these areas were more involved in learning and reward predictions than motor planning. Overall, even though the premotor signal is available across multiple brain areas, the cerebellum and ARTR contain the strongest predictive signal for decision direction. Other brain areas, such as the telencephalon, thalamus and the habenula probably regulate learning and reward prediction.

In addition, when comparing the timing of the response onset after delivery of the heat stimuli, we found that the earliest responses were detected in the telencephalon, followed by the habenula and the cerebellum (Figure 4E). Furthermore, in different trials, when the habenula responded with a longer onset time, so did the cerebellum; conversely, when the habenula responded with a shorter onset time, so did the cerebellum (Figure 4F). These overall observations (Figure 4E, F) provide certain constraints and a coarse model for how sensory information can be transformed into decision signals and motor responses: Following heat onset, the telencephalon and the thalamus receive sensory signals, with reciprocal connections between these two areas (Rink and Wullimann, 2004). This information is processed and transmitted to the habenula, where it is integrated and compared with internal states, including those representing learning and reward prediction via the dopamine and serotonin neuromodulator systems (Hikosaka, 2010; Lee et al., 2010). This modulated signal is received by the cerebellum through the mossy fibers, partly via the tegmental area (Barrot et al., 2012), together with sensory and motor signals from other pre-cerebellar nuclei (Ito, 2008), to generate the signals for motor planning.

### Preparatory neuroactivity in cerebellum and ARTR predict decision outcome on single-trial level

It is well-established that the cerebellum plays a key role in motor control and coordination (Stoodley and Schmahmann, 2010). More recent studies have suggested that it can also encode expectation for sensory outcomes (Brinke et al., 2017; Sawtell, 2017; Wagner et al., 2017) as well as that its activity is necessary to maintain cortical preparatory activity in motor planning (Gao et al., 2018). Thus, given our above observations on the pre-motor decision-dependent dPCs of the cerebellum and ARTR, we hypothesize that they may represent preparatory neuroactivity for motor response, similar to what has been reported in the cortex (Churchland et al., 2012; Gao et al., 2018; Li et al., 2016). More specifically, we asked if they would be predictive of decision outcomes.

We first looked at the trial-averaged activity of individual neurons in Cluster 4, the ipsilateral cerebellum and ARTR, and sorted the neurons according to the onset time of their activity. For correct trials, we observed a gradual increase of neuroactivity in the ipsilateral hemisphere after heat onset which continued until movement initiation, followed by a strong peak after the execution of the turn (Figure 5A (i)). In contrast, for incorrect trials, neuroactivity was reduced in general. In particular, the above gradual increase of neuroactivity was absent although the post-turn peak remained present (Figure 5A (ii)).

**Figure 5.**
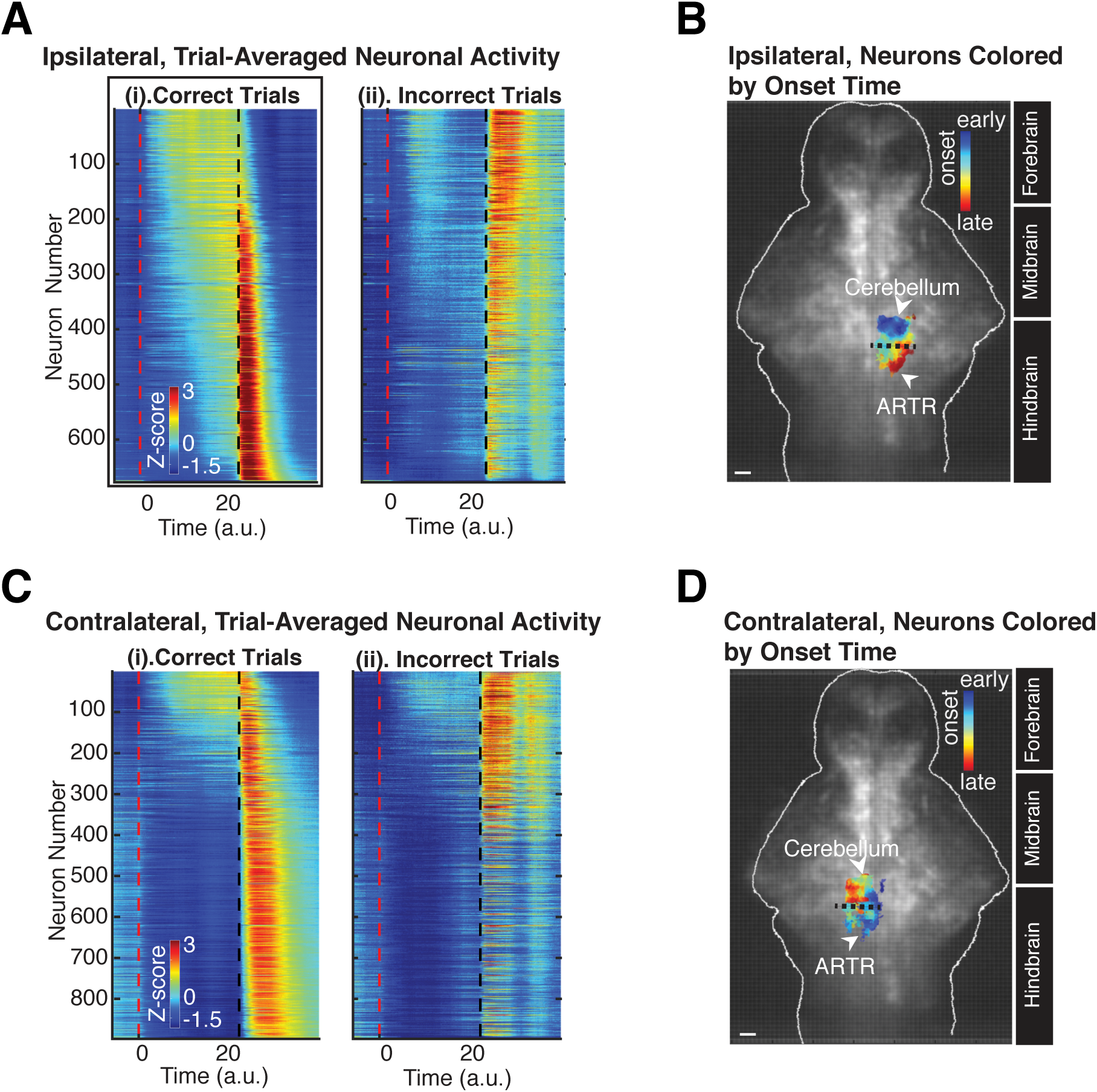
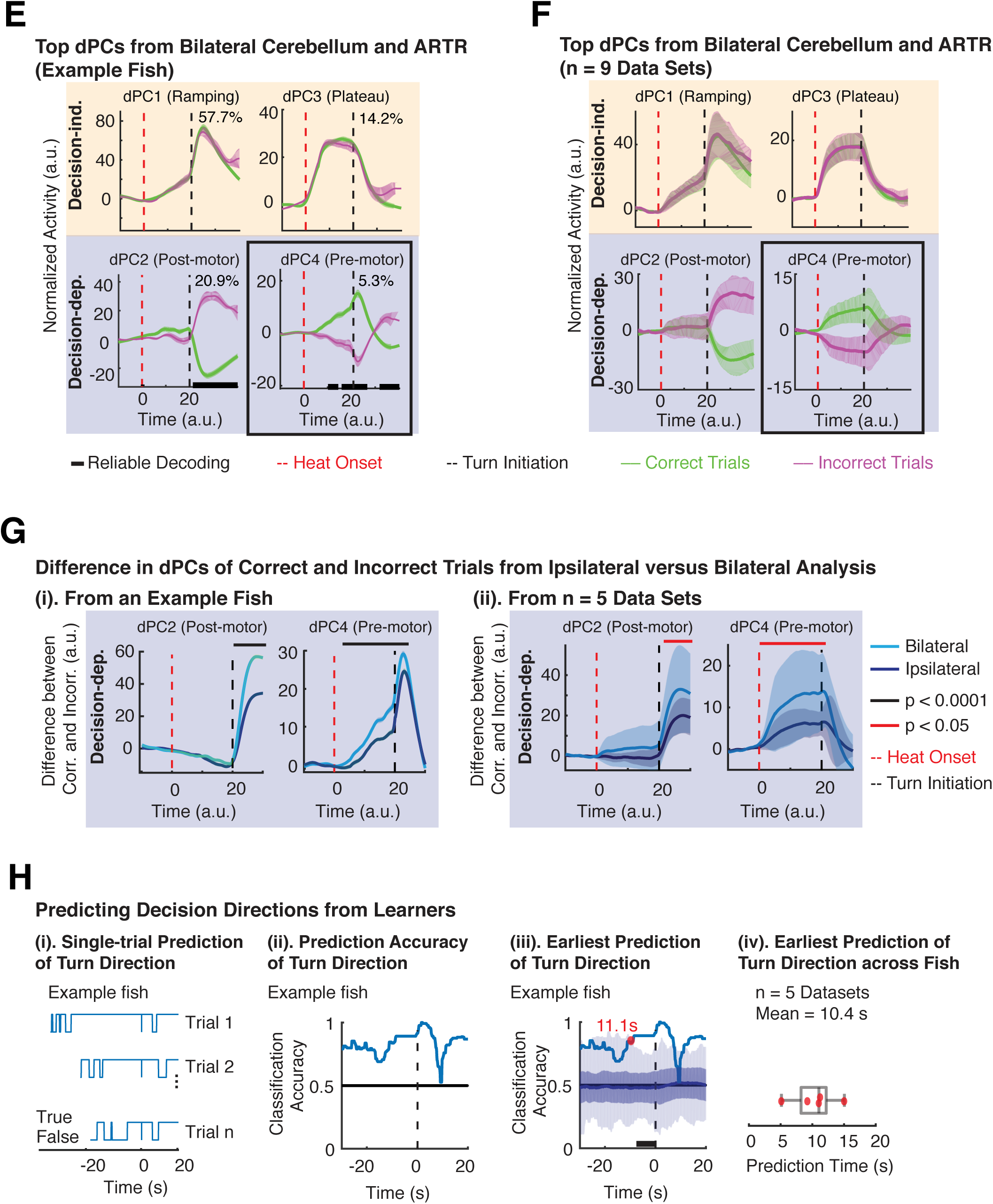
Preparatory activity from the cerebellum and ARTR predicts behavioral outcome seconds before movement initiations. (A) Preparatory activity of individual neurons in ipsilateral cerebellum and ARTR during correct and incorrect trials. Data processed to match reaction times (see STAR Methods and Figure S3A) and averaged over trials. Activity ramps up slowly in correct trials during the pre-motor period, with different onset timings. Neurons that display strong activity after turns were different in correct compared to incorrect trials. Neurons that display a strong post-turn activity in incorrect trials also have early onset time of strong pre-motor activity in correct trials. Order of neurons identical in both panels; sorted according to onset timing of preparatory activity during correct trials. Red dashed lines, heat onset; black dashed lines, turn initiation. (B) Anatomical distribution of ipsilateral neurons color-coded by onset timing of their preparatory activity. Onset time increases from anterior (cerebellum) to posterior (ARTR) in the ipsilateral hindbrain. Background: anatomical reference. Scale bar, 50 µm. (C) Pre-motor neuronal activity of the contralateral cerebellum and ARTR. Neurons are sorted by relative delay of activity during correct trials. Compared to ipsilateral side in A, this pre-motor activity is much reduced. (D) Anatomical distribution of neurons in panel (C). Scale bar, 50 µm. (E) dPCA applied to the joint neuronal activity of cerebellum and ARTR (both hemispheres). Data from one example fish. Decision-independent dPCs: dPC1 displays pre-motor ramping and post-turn peak, potentially for information integration and temporal control; dPC3, orthogonal to dPC1, displays a plateau shape during the RT, representing the heat-and-motor-evoked signals. Decision-dependent dPCs: dPC2 shows post-turn separation, which can be used as a feedback signal; dPC4, orthogonal to dPC2, demonstrates pre-motor separation, which could function as the forward signal to plan and drive action selection. (F) Averaged task-related dPCs from multiple fish (mean ± 95% CI, n = 9). The patterns of the dPCs across multiple fish are consistent with the example in panel (E). (G) Difference in dPCs of correct versus incorrect trials for ipsilateral versus bilateral population dynamics. dPCs from the bilateral (panel E, F) and ipsilateral (Cluster 4, Figure 4D) populations resemble each other, yet the difference between correct and incorrect trials is significantly increased in the joint dPCs from both hemispheres. (i) Example fish, Wilcoxon rank sum test, two-tailed. (ii) Multiple fish, Wilcoxon rank sum test, one-tailed. (H) Predicting decision directions for learner fish. (i) Examples of binary prediction at the single-trial level. (ii) Mean prediction accuracy as a function of time, obtained by aligning the binary prediction from all the trials to the movement initiation. Due to different RTs, the time points of heat onset are not aligned. Data from an example learner. (iii) Prediction time point is defined as the earliest time point (red dot) when classification accuracy (light blue curve) exceeds 99% of the shuffling tests (light shaded area). n = 100 shuffling tests; dark blue line and shaded band: mean and SD of shuffling tests. (iv) Earliest prediction times from 7 learner datasets, in 5 of which prediction precedes movement initiation (mean prediction time across these 5 animals: 10.4 s). Note that this is using non-warped time.

Sorting neurons both in correct and incorrect trials according to their activity onset time in the correct trials allowed us to draw further conclusions: We found that the identity of neurons that displayed a strong activity after turns was different in correct compared to incorrect trials (Figure 5A). Moreover, neurons that displayed a strong post-turn activity in incorrect trials were the same ones that also showed an early onset time of strong pre-motor activity in the correct trials (Figure 5A). This indicates that individual neurons displayed different ‘tunings’ during pre-motor and post-motor epochs of the task, and that pre-motor neuronal activity was not merely a subthreshold version of motor response (Churchland et al., 2010).

We then mapped the order by which neurons became active to their anatomical locations in the brain. Following an anterior to posterior gradient, we found that neurons with an earlier onset time of pre-motor activity were located in the cerebellum while neurons with a later onset of activity were mainly located in the ARTR (Figure 5B), implying a flow of information from the cerebellum to the ARTR before correct turn initiation. Moreover, the observation that the later-responding neurons in the ARTR were also those that exhibited a strong post-turn response was consistent with the recent discovery of post-turn activity in ARTR and its role to facilitate a bias in future turn direction (Dunn et al., 2016).

Compared to the ipsilateral side, the contralateral side displayed a highly reduced level of pre-motor ramping. In correct trials, the responses of the first active neurons were similar to that of the ipsilateral side, however significantly fewer neurons participated in the pre-turn ramping activity, which came to an early and abrupt halt while a similarly strong post-turn response persisted for the correct trials (Figure 5C). Subsequent active neurons which displayed a decreased pre-turn activity (Figure 5C (i)) overlapped with the contralateral cerebellum (Figure 5D). In incorrect trials, neurons that displayed a strong post-turn activity had an early onset time of pre-motor activity (Figure 5C), and mostly overlapped with the contralateral ARTR (Figure 5D).

The ARTR in each hemisphere has been shown to project inhibitory connections to the contralateral hemisphere (Dunn et al., 2016). Moreover, the parallel fibers of the cerebellar granular neurons, which feed into the Purkinje cells, span across both hemispheres of the cerebellum densely (Knogler et al., 2017). These observations provide evidence that the contralateral ARTR and cerebellum are an integral part of the learning, motor planning and action selection circuit and that the activity of the ipsilateral and the contralateral ARTR and cerebellum should be analyzed collectively. Thus, we investigated the joint dynamics of the neuronal population from Cluster 2, Cluster 4 and Cluster 5 during different task epochs. We pooled all neurons from these clusters, decomposed their joint activity into decision-independent and decision-dependent activity by applying dPCA, and focused on the top 4 dPCs, which capture >80% of the explained variance. We identified dPC1 (ramping), dPC3 (plateau), dPC2 (post-motor separation) and dPC4 (pre-motor separation) (Figure 5E), and these observations were reproducible across fish and data sets (n = 9) (Figure 5F). Although the time-dependent response of these bilateral dPCs resembled those of the ipsilateral side (Cluster 4 in Figure 4D), we found that on the quantitative level, the bilateral analysis of the circuit dynamics resulted in a significantly larger separation of the dPC trajectories during the pre-motor period for the correct and incorrect trials (87% increase, ***p < 0.0001 for example fish and 104 ± 46% (mean ± SEM), * p < 0.05 from 5 fish) (Figure 5G), consistent with the notion that two hemispheres form a joint circuit.

The observed ramping mode of dPC1 is consistent with activity that would represent a non-direction-specific, but rather a temporal aspect of decision (i.e., the decision time), while the plateau region in dPC3 is consistent with heat-and-motor-evoked signals, as the rise and fall of dPC3 coincides with heat onset and movement initiation. dPC2 represents the decision outcome after the turn—which can serve as a feedback signal for learning—while dPC4 represents the feed-forward pre-motor signal that encodes future turn direction.

In order to investigate how these neuronal activity features underlying preparation and generation of motor outputs are shaped and modulated by the learning process, we performed identical analyses in non-learners and in animals that did not undergo the ROAST assay, i.e. during spontaneous turn behavior (Figure S6). Comparing these three groups of animals provided some further hints into how these dPCs with different temporal profiles were related to different brain functions: The signals that showed decision-independent ramping and plateaus (dPC1 and dPC3) were present in a qualitatively similar fashion in both learners (Figure 5F, G) and non-learners engaged in the ROAST assay (Figure S6A, B), but were absent in fish engaged in spontaneous turn behavior (Figure S6C, D). From this we conclude that they do not encode any learning-related aspects of our task but rather, as suggested before, are related to the representations of the heat stimulus and the timing of the movement initiation, respectively. In contrast, the signal predictive of the turn direction (dPC4) was only observed in learners (Figure 5E, F); this component was not predictive in non-learners and was absent during spontaneous turns (Figure S6E, F). These observations imply that this component was directly associated with the animals’ ability to learn, plan and execute their turns in the target direction. This interpretation and the absence of dPC1 and dPC4 in animals performing spontaneous turns is also consistent with the fact that in absence of the ROAST paradigm, they both do not bear any behavioral relevance: During spontaneous behavior, there is no urgency, nor a defined, task-related motivation for the animal to initiate turns in a specific direction. Finally, since the post-turn feedback signal (dPC2) appeared in all three groups (Figure 5E, F, S6A-D), we conclude that it represents an isolated downstream circuit of the cerebellum, likely arising from the inputs of the inferior olive (Marr, 1969).

Next, given our observations on the pre-motor decision-dependent dPCs in the learner fish, we set out to quantify how well these predicted the outcome of the turn before movement initiation at the single-trial level. Using a linear binary decoder (Kobak et al., 2016) (see also Figure S4C), we first predicted the turn direction at the single-trial level as a function of time while aligning all trials based on their movement initiation time point (Figure 5H (i), Movie S3). Subsequently, we calculated the time-dependent classification accuracy (Figure 5H (ii)) and defined the prediction time as the earliest time point at which the turn direction was able to be decoded from the decision-dependent dPCs reliably (i.e., when actual classification accuracy exceeded the accuracy of 99% of the shuffled tests in at least ten consecutive time points, see STAR Methods). Figure 5H (iii) shows the prediction time in our example learner dataset, for which we found a value of 11.1 seconds; non-learners and spontaneous fish were not predictive (Figure S6E, F). This result was reproducible across multiple learners (n = 5) for which we found an average prediction time of 10.4 seconds (Figure 5H (iv)). This time scale is different from the millisecond time scale that has been traditionally associated with the role of the cerebellum in motor control (Buhusi and Meck, 2005), but well within the range of interval timing (i.e. seconds to minutes) involved in decision making and working memory (Buhusi and Meck, 2005; Tetzlaff et al., 2012). Taken together, the difference in the cerebellar pre-motor activity between learners, non-learners and spontaneously behaving fish showed that the preparatory activity for planning turn directions is tightly linked with an animal’s learning ability, and the time course of the preparatory activity provides further evidence for the involvement of the cerebellum in motor planning.

### Ipsi-contralateral difference in neuroactivity in the cerebellum and ARTR drives turn direction on single-trial level

It has been shown that turn direction during spontaneous behavior in freely moving larval zebrafish is biased in the direction of the previous turn by the ipsilateral activity of the ARTR, and that ARTRs in the two hemispheres have mutually inhibitory connectivity (Dunn et al., 2016). This bilateral ARTR network provides the basic circuity necessary to allow for generation of mutually exclusive decision outcomes by two competing neuronal populations (Gold and Shadlen, 2007; Wang, 2012; Wong et al., 2007). We speculated that the decision for the turn direction might arise from an ipsilateral-contralateral interaction between the neuroactivity in the cerebellum, with the winning side determining the decision outcome, and asked if this explained the observed turn directions at the single-trial level.

We tested this idea by comparing the difference in the neuronal activity of the cerebellum and ARTR between the two hemispheres during the pre-motor period. After applying dPCA on neurons from both hemispheres, we reconstructed and mapped the neuronal signal only from the pre-motor decision-dependent dPC4 back to the neuronal time series, using the linear encoder that we had obtained by applying dPCA (Figure S7A and STAR Methods). Compared to the original neuronal time series, this reconstructed time series only contained pre-motor decision-dependent information but had the same dimension (i.e. neurons × time). Thereby, we managed to preserve the information on the location of each neuron and compare the ipsi-contra difference in activity. An example of the ipsi-contra difference of reconstructed neuronal activity as a function of time is shown in the Figure 6A. As can be seen, this difference in pre-motor activity is higher during correct trials, and lower during incorrect trials, in the ipsilateral than the contralateral side. In order to compare the pre-motor activity only and avoid contamination from post-motor activity, we evaluated the reconstructed difference in neuronal activity between the ipsilateral and contralateral hemispheres only during the pre-motor period from −4 to −2 seconds before turn initiations (see STAR Methods).

**Figure 6.**
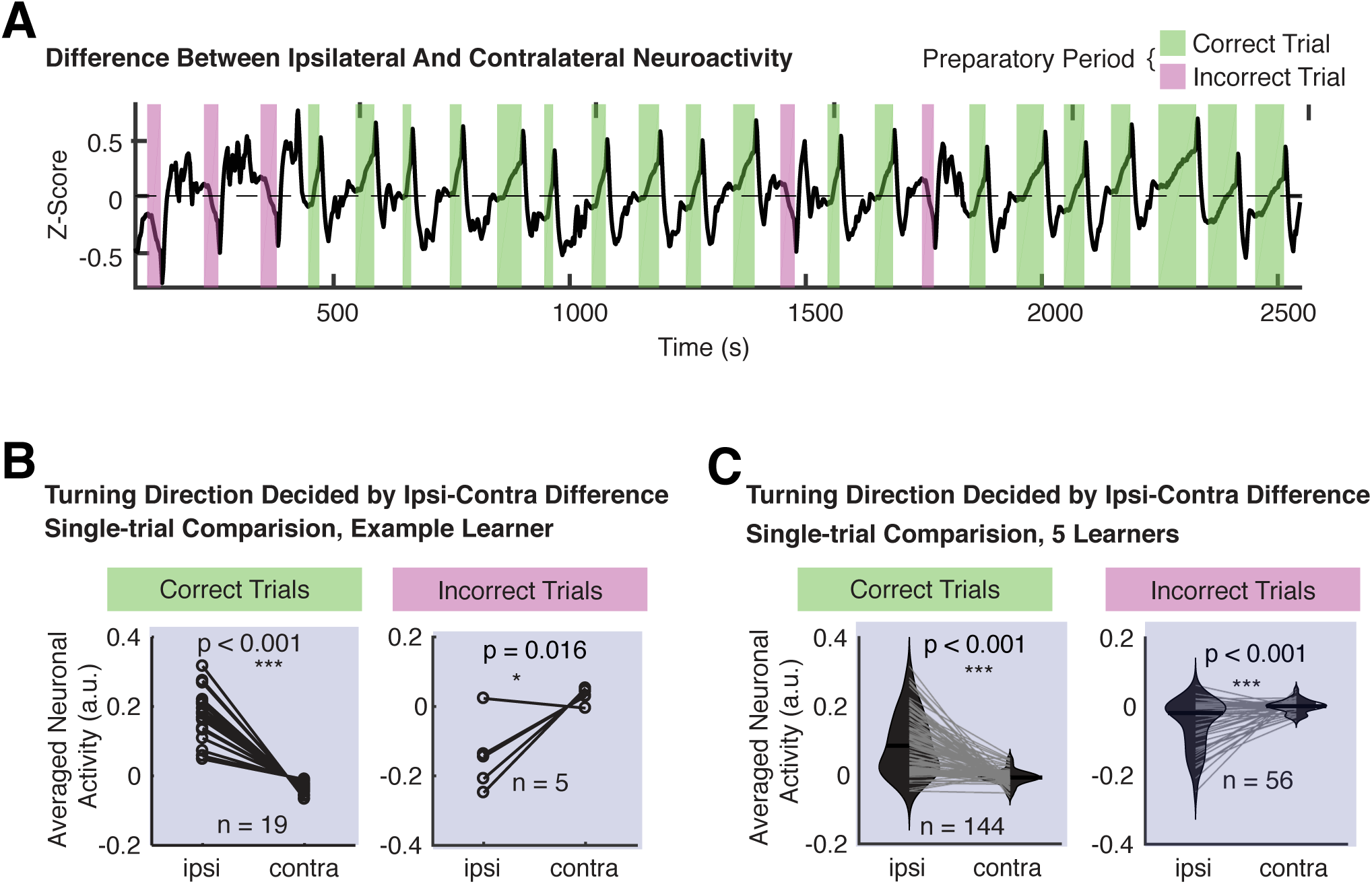
Turning direction explained by ipsilateral-contralateral competition in the cerebellum and ARTR. (A) Difference between reconstructed neuronal activity from the ipsilateral and contralateral cerebellum and ARTR as a function of time. See Figure S7A for the reconstruction process from dPCs. Data is the pre-motor decision-dependent reconstructed activity from an example learner. The neuronal activity is stronger in the ipsilateral side during correct trials (green shading) and weaker during incorrect trials (magenta shading). (B) Comparison of pre-motor reconstructed neuronal activity between ipsilateral and contralateral sides, shown trial-by-trial, from an example learner. Pre-motor activity is averaged from 4 to 2 s before turn initiation. For the learner, the neuronal activity in the ipsilateral side is significantly higher than the contralateral side in the correct trials, and significantly lower in the incorrect trials, on a trial-by-trial basis. Wilcoxon signed rank test, one-tailed. (C) Comparison of the ipsilateral-contralateral difference at the single-trial level from data pooled across multiple learners. In the 144 correct trials from 5 learners, the ipsilateral pre-motor activity (median > 0) is significantly higher than the contralateral side (***p < 0.001), while in the 56 incorrect trials, the ipsilateral pre-motor activity (median < 0) is significantly lower than the contralateral side (***p < 0.001, left panel). Wilcoxon signed rank test, one-tailed.

At the level of individual learners, we found a significantly higher level of preparatory activity in the ipsilateral side than the contralateral side in correct trials (19 trials), but a significantly lower level of activity in the incorrect trials (5 trials) (Figure 6B). In contrast, at the level of individual non-learners, no significant difference between the activity on the ipisi-and the contralateral side was observed, both in correct and incorrect trials (Figure S7B). These observations were consistent across fish and when all trials were pooled (Figure 6C & S7C). We only observed this mirroring relationship between the ipsilateral and contralateral side in dPC4 across animals, and not any of the other three top dPCs. This inverse relationship between the neuroactivity of ipsilateral and contralateral side was only reflected by the dPC4 and not any of the other top three dPCs. These observations support our idea that cross-hemisphere difference of neuroactivity in the cerebellum and ARTR during the motor planning period forms the basis of the decision, with the winning side determining the decision outcome on the single-trial level. This idea is also consistent with existing decision-making frameworks based on attractor or diffusion models (Shadlen and Kiani, 2013; Wang, 2012) of competing neuronal populations or neural signals.

### Single-trial reaction time is predicted by the ramping rate of the joint population dynamics in the cerebellum and ARTR

A number of previous studies have shown that in various decision-making tasks, the behavioral reaction time (RT), i.e. the time from the onset of the stimulus or go cue to movement initiation, is linked to the velocity of the increase of cortical preparatory activity (Afshar et al., 2011; Hanes and Schall, 1996; Murakami et al., 2014; Shenoy et al., 2013; Wang, 2012). Since we observed the ramping signal in the decision-independent pre-motor component of the neuroactivity in cerebellum and ARTR (Figure 5G, dPC1), we asked if the velocity of this ramping activity, i.e. the ramping rate of dPC1, would be correlated with RT at the single-trial level.

Our behavioral paradigm lacks a go cue and movements were self-initiated by the animals, which results in variability of RTs across trials (Figure 1E, S1E). Thus, we expected that a potential fixed relationship between the speed of ramping and the RT would be observable by comparing trials with different RTs. In doing so we indeed found trials with a shorter RT to exhibit a higher rate of ramping and those with a longer RT a lower rate of ramping activity (Figure 7A). This was more clearly seen by grouping all trials in four bins of RTs for which a quasi-linear relationship between RT and ramping rate was observed (Figure 7B).

**Figure 7.**
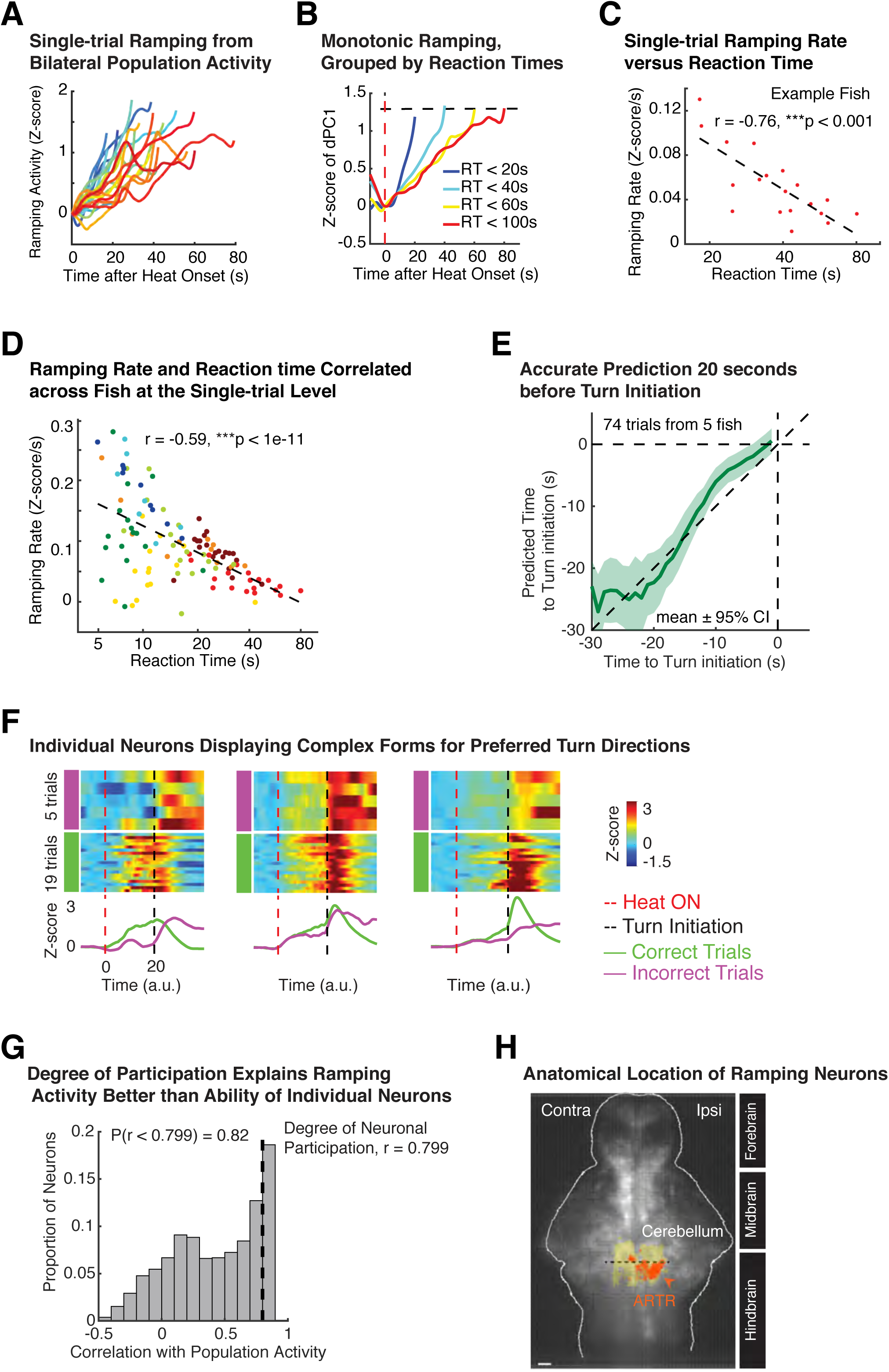
Ramping rate of the population dynamics quantitatively explains reaction time (RT). (A) Examples of ramping activity (as Z-score of dPC1) for individual trials after heat onset. Data from the bilateral cerebellum and ARTR (with explained variance > 50%) of a learner. Different trials are color-coded. (B) Quasilinear monotonically ramping population activity reaching a common threshold independent of RTs. Population activity (dPC1), grouped as 4 categories of RTs and then averaged across trials. Same data as panel (A). Each trace is from heat onset (red dashed line) to the first turn. Different categories are color-coded by their corresponding RTs. Short RT, such as the blue trace, corresponds to a steeper slope. Black dashed line indicates arbitrarily chosen threshold. (C) Ramping rate of population dynamics (dPC1) as a function of the RTs at the single-trial level. Ramping rate is calculated by linear fit of pre-motor ramping activity (from heat onset to 1 s before turn initiation) versus time (see Figure S8A). Data points are single trials. Linear regression shows a strong negative correlation between the ramping rate and the logarithm of the RT. Data from an example learner. (D) Ramping rate of population dynamics from the cerebellum and ARTR is inversely correlated with the RTs at the single-trial level across multiple animals. Data points are single trials. 8 training blocks in different colors from 5 animals, total 114 trials. Despite individual variance in learning, RTs and neural activity, the inverse correlation between ramping rate and RT is highly robust. (E) Increase of prediction precision for movement initiation time as actual initiation time approaches. Prediction is made using linear fit (see Figure S8D for parameter estimation) at the single-trial level (see also Figure S8E). Predicted values (mean ± 95% CI) are plotted against the actual values. Data from 5 fish. (F) Three example neurons to illustrate the complex forms of pre-motor and post-motor activity across trials, and more importantly pre-motor ramping is not present in these neurons at the single-trial level. Warped time series (See Figure S3) are used to show and compare multiple trials. (G) Distribution of correlation between activity from single neurons and population ramping activity. 82% of the neurons show correlation <0.799, hence only 18% of neurons exceed the correlation between neuronal participation ratio and population activity. In other words, ramping activity is better explained by the level of neuronal participation than by individual neurons. (H) Anatomical location of the neurons (top 18% in (I)) whose correlation coefficients to population ramping activity exceed the correlation between degree of neuronal participation and population activity. These neurons mainly overlap with the ipsilateral ARTR, colored red. Other active neurons from the rest of the cerebellum and ARTR are indicated in yellow. Scale bar, 50 µm.

We then calculated the ramping rate by a log-linear fit of the population activity (Z-score of dPC1) from heat onset to one second before the turn (Figure S8A) and found the ramping rate to be linearly correlated with the logarithm of the RT at the single-trial level (R = −0.76, ***p < 0.001, Figure 7C). We further confirmed that such an inverse correlation between ramping rate and RT was indeed present across animals at the single-trial level (R = −0.59, ***p < 1e-11, 8 training blocks from 5 animals, 114 trials in total) while when RTs were averaged across trials of each fish, systematic differences in the average RT of each fish were observed (Figure 7D). This inverse relationship between RT and ramping rate was also found in the correct trials of non-learner fish (Figure S8B, C), suggesting that the ramping activity was only task-dependent but not tightly linked to learning performance of the animal. In other words, the signal for when to move was dissociable from the signal for which direction to move to.

Given, the above inverse correlation between ramping activity and RT, we asked if it would allow us to predict the time for turn initiations on the single-trial level. Using log-linear regression, we obtained the fit parameters for a linear ramping (log10(*RT*) = −3.94 \**Ramping Rate* + 1.66) for the learner and non-learner fish (Figure S8D) and used this linear relationship to make a prediction about the expected turn initiation time at the single-trial level at any given time point *t* after the heat onset. To do so, we calculated the ramping rate at each time point *t* in a given trial by using the information on the past activity in that trial and used the obtained ramping rate at time point *t* to calculate the RT using our above linear regression (Figure S8D) and subsequently the predicted turn initiation time point by using *RT* – *t* (Figure S8E).

As expected, we found the predicted turn initiation time to display an increased accuracy as *t* approached the turn initiation time point and to be highly predictive for actual turning time within a 20-second window before the turn (22.3 s ± 2.5 s when *t* = 20 s, mean ± SEM), at the single-trial level (Figure 7E, Movie S3). In principle, such ramping could be facilitated by either a gradual increase of activity of individual neurons, which seemed less likely as individual neurons displayed a complex form at the single-trial level (Figure 7F), or from more neurons joining the population of active neurons over time and participating in the generation of ramping activity on a population level. We proceeded to differentiate between these two possibilities.

To investigate the degree at which an increasing pool of active neurons was the driver of the observed ramping activity, we first classified the states of activity of each neuron at a given time point in a binary fashion as “active” or “inactive” depending on whether its activity was above or below its own standard deviation (SD). We then determined the degree of neuronal participation by calculating the fraction of active neurons at each time point and compared this to the population activity, as reported by dPC1, as a function of time. We compared this to the scenario in which the ramping activity would be explained by an increasing level of activity of individual neurons. To do so, we used linear correlation to measure the similarity of activity of all individual neurons to the population ramping during the pre-motor period, obtained the correlation coefficients for each neuron, calculated the distribution of these correlations and compared this distribution to the correlation obtained between the neuronal participation and the population ramping.

By using linear correlation analysis on the population level, a high level of similarity between the ramping activity (dPC1) and the degree of neuronal participation was observed (r = 0.799, ***p < 0.0001). In contrast, the observed correlation of single-neuron activities with the population ramping activity was lower and ∼82% of all neurons showed a lower degree of correlation with the population ramping activity compared to the degree of neuronal participation (Figure 7G). The remaining 18% of neurons overlapped with the ipsilateral ARTR (Figure 7H), which presumably receive collective ramping signals from the cerebellar neuronal populations.

Pinpointing the source of the above ramping activity on the single-trial level requires further and more detailed investigations. However, two observations in our data hint that the habenula is an important player: First, the population dynamics of the habenula displayed ramping (Figure 4E). Second, neuroactivity after heat onset had an earlier start in the habenula than in the cerebellum (Figure 4E) and the timings were correlated between these two areas (Figure 4F). Also, given its known role in learning and generation of reward signals (Hikosaka, 2010; Matsumoto and Hikosaka, 2007), the habenula is thus very likely a key region providing inputs to the cerebellum, potentially via the tegmental area (Barrot et al., 2012; Quina et al., 2015). These points suggest that that the observed ramping activity in the cerebellum and ARTR is unlikely to be exclusively self-generated.

## DISCUSSION

Decision making and generation of goal-directed behavior relies on coordination and integration of brain functions such as the processing of sensory information, memory, motivation or internal states, predicting reward and coordinating motor actions, all of which typically reside in vastly separated brain regions (Abbott et al., 2017). Studies on decision making however have been mainly focused on rodents or primates in which, due to technical limitations, recording of neuroactivity at high spatial and temporal resolution during decision making tasks can only be performed in parts of the brain, typically within the cortex (Gold and Shadlen, 2007; Murakami and Mainen, 2015; Svoboda and Li, 2018). Nevertheless, a hallmark of motor planning, the preparatory neuroactivity, has also been reported in different subcortical areas (Guo et al., 2017; Horwitz and Newsome, 1999; Kunimatsu et al., 2018; Liu et al., 2013; Svoboda and Li, 2018).

Supported by the above conjecture on brain-wide processing of information for goal-directed behavior, in this work we have used an operant learning task to investigate the neural basis underlying decision making in larval zebrafish. The combination of our behavioral assay with high-speed whole-brain calcium imaging based on LFM and advanced data analysis has allowed us to observe global and local brain state dynamics at single-neuron resolution, and their relationship to behavioral and stimulus features at the level of individual trials. We have found that on the global level, brain states display stimulus-triggered recurrence and a decision-related bifurcation at the trial-by-trial level, in which pre-turn neuroactivity distributed across the brain is predictive of the future turn direction. On the level of brain regions, we identified predictive neuroactivity for decision direction that are spatially distributed across multiple brain areas but also highly concentrated in the cerebellum (Figure 4D & E, S5D). Based on the observed neuroactivity in different brain areas and their known connectivity, we proposed a model for how sensory information was transformed into decision and motor signal, and how learning and reward prediction was facilitated in this process (Figure 8). The telencephalon (Portavella and Vargas, 2005), the thalamus (Komura et al., 2001; Zhu et al., 2018) and the habenula (Hikosaka, 2010; Lee et al., 2010; Matsumoto and Hikosaka, 2007) are known to be involved in the regulation of learning and reward prediction, and can instruct the cerebellum on action selection (Figure 8). Upon heat onset the sensory signal is relayed to the telencephalon (Cluster 1 in Figure 4D) which on one hand suppresses neuroactivity in the thalamus (Cluster 6 in Figure 4D) and on the other hand activates the habenula (Cluster 3 in Figure 4D). The habenula displays a ramping-like activity during the pre-motor period. This activity is fed to the cerebellum, in part via the tegmentum though mossy fibers, and is consistent with a signal encoding for reward prediction. Our assay involves a learning and motor planning component. Thus, in contrast to heat-evoked sensorimotor transformation which involves a hindbrain local circuit (Haesemeyer et al., 2018), the overall functional circuitry spans across the entire brain (Figure 8), even though both studies observed strong response to heat in the telencephalon and the habenula (Figure 4).

**Figure 8.**
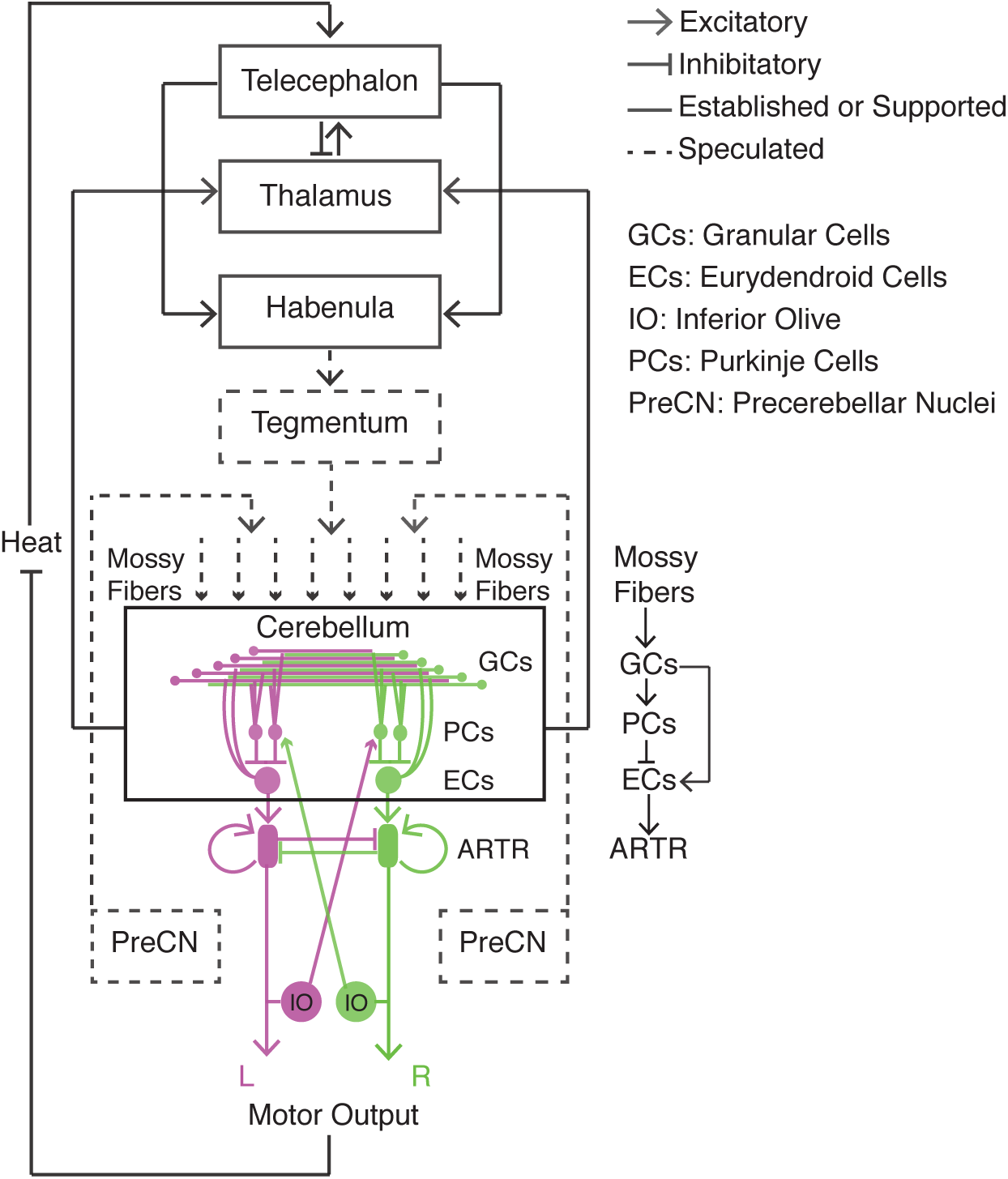
Proposed model for decision-making network in cerebellum and ARTR. Schematic overview of the neurocircuitry underlying sensory information flow, learning, motor planning and action selection in our task: The telencephalon, thalamus and habenula regulate learning and reward prediction, and instruct the cerebellum on action selection. The cerebellum integrates multimodal information, prepares and generates movements. Following movement initiation, the cerebellum sends motor information back to the thalamus, thereby forming a closed loop for reinforcement learning. After heat onset, the sensory signal is relayed to the telencephalon which suppresses neuroactivity in the thalamus and activates the habenula. The habenula displays a ramping activity during the pre-motor period, encoding for reward prediction, feeds into cerebellum, in part via the tegmentum, though mossy fibers. The input to the cerebellum is further amplified through the divergence circuitry on the granular cell level and subsequently integrated at the Purkinje cells. Then the Purkinje cells modulate the output of the eurydendroid neurons via inhibitory connections which feed into the ARTR. At the same time, the granular cells send excitatory input into the eurydendroid neurons, instructing the downstream ipsilateral ARTR. Following movement initiation, the inferior olive (IO) feeds back the signal of the motor output to the contralateral Purkinje cells via the climbing fibers and subsequently the motor information in the cerebellum is sent to the thalamus and the telencephalon.

This input to the cerebellum could be further amplified by the divergent mapping of the mossy fiber input to the granular cells (Huang et al., 2013; Ito, 2006; Li et al., 2013) and then integrated at the Purkinje cell layer, which also receives excitatory input from the contralateral inferior olive (IO) through the climbing fibers (Ito, 2008). Purkinje cells modulate via inhibitory connections the output of the eurydendroid neurons which at the same time also receive direct excitatory connections from the granular cells (Ito, 2008) and feed into the ARTR, i.e. the hindbrain oscillator (Figure 8). The underlying divergence of this circuitry at the level of the granular cells and their abundance in the cerebellum (Ito, 2006) explains on one hand our observation on the neuronal participation model (Figure 8) and accounts for the granular cells as the main source of the observed ramping On the other hand, the observed contribution to the ramping by a small neuronal population with a gradual increase of activity (Figure 7G) can be explained by the activity of the ARTR neurons, which presumably receive input from the eurydendroid neurons. This notion is also directly supported by the observation that this small neuronal population is anatomically overlapping with the ARTR neurons (Fig 7H).

In our study, the observation of preparatory activity in the cerebellum and ARTR provide support for two key aspects of decision making: the decision outcomes and decision timing (see also Movie S3). The bilateral relationship between ARTR and the cerebellum can be viewed as a ‘competition-cooperation’: During the pre-turn period, the turn direction is driven by the differential activity between the two sides resulting in a turn to the side with the higher level of activity. The timing of the movement initiation on the other hand is determined by the ramping rate of the joint population activity from both hemispheres. After motor response, two feedback signals are sent: The IO activates the Purkinje cells via climbing fibers in the contralateral side of the cerebellum (Apps and Garwicz, 2005), while a self-excitatory ipsilateral feedback is sent to the ARTR (Dunn et al., 2016). The signal to the contralateral Purkinje cells can serve as an error signal that, together with inputs from other brain areas, facilitates synaptic plasticity (Albus, 1971; Apps and Garwicz, 2005; Dempsey and Sawtell, 2016; Marr, 1969).

Previous models of decision making based on ramping-to-threshold have established a link between the activity of two competing neuronal populations and their corresponding selectivity for decision outcomes and decision times (Gold and Shadlen, 2007; Shadlen and Kiani, 2013; Uchida et al., 2006; Wang, 2012). Supported by our observations, the above competition-cooperation model—in which the joint bilateral (as opposed to the unilateral) activity determines the timing of the decision, while the differential activity drives the decision outcome—provides evidence that competition and cooperation of neuronal populations can exist at the same time, in the same brain region and within the same behavior. This is distinct from a previously proposed model in which temporal transitions in patterns of cooperation and competition between neural systems are driven by context and task (Cocchi et al., 2013). This new model provides new insights into the dynamical interactions among neuronal populations for further studies on computational models of decision making, especially given the fact that large-scale single-cell-resolved imaging at high speed, subthreshold neuronal activity recording via voltage indicators, and determination of the connectome between cerebellar neurons should soon become feasible (Hildebrand et al., 2017; Miyazawa et al., 2018; Prevedel et al., 2016).

The timescales of the cerebellar preparatory activity reported in the present study are more consistent with the timescales of cognitive process, such as decision making, rather than the timescales of control: Previous studies of motor control in the cerebellum found the onset neural activity to take place hundreds of milliseconds before onset of movement (Johansson et al., 2016). In comparison, the cerebellar preparatory activity underlying the timing of turn initiation and decided direction which we describe in the present study unfolds on a longer timescale (∼10 s). Such a prolonged time scale is task-specific (Waskom and Kiani, 2018) and a feature of our behavioral paradigm. This is because first, the aversive heat stimulus in this task is mild and thus imposes no immediate urgency for fish to react, until the steady state temperature represents an aversive stimulus for the animal. Second, our task has a learning component, the direct relevance of which for the involved longer timescales is shown by comparison to non-learners and fish undergoing spontaneous movements in which the predictions of turn initiation and direction are not possible to the same degree (Figure S6). Therefore, the reported time scales here represent those involved in preparation of motor response based on cerebellar learning and are more consistent with what has been reported in the literature in connection to tasks that require working memory (Howe et al., 2013; Kunimatsu et al., 2018; Soon et al., 2008; Tetzlaff et al., 2012), rather than what has been associated with motor control (Buhusi and Meck, 2005).

In mammals, the neuronal processes underlying decision making and preparation of goal-directed movements has been mainly attributed to the cortex (Miller, 2000; Shenoy et al., 2013; Svoboda and Li, 2018), while motor coordination functions, such as fine motor control and execution have been traditionally associated with the cerebellum (Glickstein, 1993; Manto et al., 2012). In this study, we have shown that the cerebellum can also be involved in higher level brain functions, the characteristic time scales of which are significantly longer. The observed bi-lateral differential population activity during the pre-turn period on the one hand provides a highly predictive signal for future turn directions; On the other hand, the joint ramping of the same neuronal population encodes for the timing of turn initiation. Moreover, we found complex patterns of tuning to different task epochs at the single-neuron level. All of these observations are reminiscent of previously reported cortical preparatory activity during decision making tasks in mammalian brains and thus ascribe to the cerebellum, as a major contributor in decision making, a more cognitive function than conventionally assumed. Our observations and interpretations are consistent with a number of recent studies which have challenged the traditional views of the cerebellum by providing evidence for cognitive functions such as in learning, language and reward prediction (Buckner, 2013; Giovannucci et al., 2017; Raymond and Medina, 2018; Sokolov et al., 2017; Strick et al., 2009; Timmann and Daum, 2007).

Further support for the general idea that the cerebellum might be involved in a broader range of cognitive functions comes from three unrelated lines of observation: First, evolutionarily, the cerebellum is a highly conserved brain structure with, after neocortex, the highest rate of growth during evolution (Finlay and Darlington, 1995). At the same time, the ratio of the number of neurons between neocortex and the cerebellum has remained relatively constant (Herculano-Houzel, 2012). Consistent with these observations, human functional brain imaging has revealed functional connectivity between the majority of the cerebellum and the corresponding association cortex, including the prefrontal cortex (Buckner, 2013). All of these findings imply a hub-like function of the cerebellum and a high degree of cerebellum-mediated intercortical connectivity. Finally, a very recent study has provided the first direct evidence that the cerebellum is necessary to maintain the cortical preparatory activity, and vice versa, and that disrupting this cortico-cerebellar loop decreases the correct rate of behavioral response, i.e. decision direction, as well as the reaction time without affecting the motor execution (Gao et al., 2018).

We conclude by pointing out three open questions that emerge from the present study: First, how do the cerebellar granule cells, via the mossy fibers, use and process various sensory, learning and reward related signals, to facilitate motor planning? Based on the highly concentrated and predictive information in the cerebellum, we propose that the cerebellum acts as an integrator and amplifier. It would be important to understand on the single cell level how individual granule cells integrate and encode the different streams of information relevant to the circuit (Giovannucci et al., 2017). This can be done by testing the causal relationships between other brain areas and the cerebellar granule cells. Second, what are the microcircuits and wiring diagrams within the cerebellum and between cerebellum and ARTR, and how do these microcircuits interact with global brain states? In the present study, the activities of the Purkinje, granular and eurydendroid neurons were not explicitly differentiated, even though our results suggest that the post-turn activity arises from the Purkinje neurons, while the forward ramping signal results from the population dynamics of granular cells. Answers to this question would require reconstruction of the connectome, faster neural indicators, and specific transgenic lines. It will provide new insights into the principles and implementation of neural algorithms. Third, what are the implications of our observations on the role of the cerebellum in motor planning and action selection in other species, in particular mammals? Recently, it has been shown that the cerebellar-cortical loop is necessary to maintain the preparatory activity during motor planning (Gao et al., 2018). Our work expands on this observation by demonstrating the generality of the relevance on cerebellar neurodynamics in vertebrates and by quantitatively demonstrating how decision time and decision outcome could be decoded from this activity. Given current advances in large-scale, volumetric and high-speed calcium imaging (Ji et al., 2016; Yang and Yuste, 2017; Weisenburger and Vaziri, 2018), behavioral studies similar to ours can be expected to become possible in rodents, and to allow the activity of the entire cerebellum to be recorded. On a fundamental level, understanding how cerebellar activity is related to the timing and outcomes of action selection can provide insights into the functional role of the cerebellum during evolution, and the precise manner of its co-evolution with the cortex.

## AUTHOR CONTRIBUTIONS

Q.L. performed experiments, contributed to the design of data analysis approach, analyzed data and wrote the manuscript. M.H. built, tested and characterized behavioral setup for ROAST with technical support from F.E. F. Schlumm performed spontaneous behavior experiments. ROAST was originally designed and built by J.L. and D.R. under guidance by F.E. and A.S. T.N. built microscopy platform and image reconstruction pipeline and contributed the final version of the manuscript, A.V. conceived and led the project, designed experiments and the data analysis approach and wrote the manuscript.

## ACKNOWLEDGEMENTS

We thank J. Manley, O. Skocek and C. Kirst for discussions on data analysis. We thank C. Bargmann, G. Maimon, M. E. Hatten, J. Manley for suggestions on the project and comments on the manuscript. We thank A. Kaczynska, A. Afolalu and J. Hudspeth for housing zebrafish. Q.L. has been partially supported by a Leon Levy Fellowship. T.N. has been supported by a Leon Levy Fellowship and a Kavli Fellowship. This work was supported in part through funding from the US National Science Fundation (NSF) awards DBI-1707408, PHY-1748958, the Intelligence Advanced Research Projects Activity (IARPA) via Department of Interior/Interior Business Center (DoI/IBC) contract number D16PC00002, National Institutes of Health (NIH) Grant R25GM067110, the Kavli Foundation, and the Gordon and Betty Moore Foundation Grant No. 2919.01.

## STAR METHODS

### KEY RESOURCES TABLE

| REAGENT or RESOURCE | SOURCE | IDENTIFIER |
| --- | --- | --- |
| Experimental Models: Organisms/Strains |  |  |
| Tg(*elavl3:H2B-GCaMP6s*) | (Vladimirov et al., 2014) | N/A |
| Software and Algorithms |  |  |
| µManager | Open Imaging | https://micro-manager.org/ |
| MATLAB (Behavioral and imaging analysis, modeling) | MathWorks | https://www.mathworks.com/products/matlab.html |
| FIJI (ImageJ) (Image Processing) | NIH | http://fiji.sc |
| Demixed Principal Component Analysis (dPCA) | (Kobak et al., 2016) | https://github.com/machenslab/dPCA |
| t-Distributed Stochastic Neighbor Embedding (t-SNE) | (Maaten and Hinton, 2008) | https://lvdmaaten.github.io/tsne/ |

### CONTACT FOR REAGENT AND RESOURCE SHARING

Further information and requests for resources and reagents should be directed to the lead contact, Alipasha Vaziri.

### EXPERIMENTAL MODEL AND SUBJECT DETAILS

#### Animal subjects

Experiments were carried out in accordance with protocols approved by the Institutional Animal Care and Use Committee. Zebrafish (*Danio rerio*) lines used in this study for imaging and behavioral experiments were 7-8 day-post-fertilization *Tg(elavl3:H2B-GCaMP6s)* (Vladimirov et al., 2014) in Nacre or Casper mutant background. Adult fish were housed in a facility at 28.5 °C with lights on between 8 am and 10 pm. No statistical methods were used to pre-determine sample size.

### METHOD DETAILS

#### Whole-brain Calcium Imaging with the Operant Conditioning Task

7-8 day-post-fertilization zebrafish larvae were embedded in 2% low-melting-temperature agarose in a custom-made chamber on a glass slide and then immersed in fish water. The agarose around the tail, caudal to the swim bladder, was removed to free the tail of the larvae and then incubated for 6-8 hours for further solidification. Under this head-fixed condition, larvae were placed above a camera (Grasshopper3, PointGrey) for tracking tail movements at 160 Hz and placed under a 20×/0.5-NA water-immersion objective (Olympus) for whole-brain calcium imaging with an upright light-field microscope at 10 Hz (Nöbauer et al., 2017; Prevedel et al., 2014). The tail was illuminated with a near-infrared (NIR) 950 nm light-emitting diode (LED), and GCaMP was excited with a blue LED (pe-2, CoolLED). Custom-written MATLAB (MathWorks) software was used to extract the tail angle, using the resting tail position as the reference, with positive angles representing right turns and 0 for a straight tail. Heat stimulus was delivered using 980-nm fiber-coupled laser (Roithner Lasertechnik), collimated via a collimator (Thorlabs, F220FC-1064) to a small beam (measured power: 300 mW) and then projected to the head of the fish along the midline, with the spot diameter of 2.0 mm. A 950-nm bandpass filter (FB950-10, Thorlabs) was placed in front of the Grasshopper camera to reject scattered visible light and cross-talk from the heat stimulus. Real-time heat stimuli were controlled by a data acquisition board (USB-6008, National Instruments) and modulated by the extracted tail angle exceeding a threshold (35°) in a closed loop. For movement events with multiple tail deflections, only the first deflection was considered.

Image acquisition of the LFM was controlled using Micro-Manager (Stuurman et al., 2007) and triggered by the custom MATLAB software. The whole setup was controlled from a dual-CPU workstation (Z820, HP). 3D reconstruction of the LFM images was carried out offline with the volume of ∼700 μm × 700 μm × 200 μm with 51 z-planes and fed into a custom-written pipeline for neuronal signal extraction (Mukamel et al., 2009; Prevedel et al., 2014).

#### Training Protocol and Animal Behavior

Typically, each recording consisted of 20-25 trials and lasted for one hour. In each trial, heat stimulus was delivered 5 seconds after the trial started and was terminated immediately when a turn in the correct direction and exceeding a threshold (35 degree) was detected. If an animal failed to make a correct movement within 100 s, the laser was switched off and the fish received a 20 s break before the start of the next trial. The full training protocol for each fish consisted of two training blocks (one left- and one right-training block), with each block containing 20-25 trials. Since individual fish exhibited a bias for a specific direction (Li, 2013), the reward direction in the first training block was chosen against this bias and was subsequently reversed in the second block. Prior to the first block, 3-5 probe trials were conducted to determine the pre-existing bias of individual fish. For learners, the same larvae were imaged twice for two training blocks, over a total of two hours, and with a 0.5 h break in between. A given trial was classified as correct when the first heat-evoked turn of the animal was in the reward direction, and a fish was defined as a “learner” when the asymptote of the learning curve (modelled by a sigmoidal function that was fit to the outcomes) in both blocks reached the threshold of 70% correct. Fish that learned in the first but not in the second block were defined as intermediate learners and their data was not included in the data analysis. If a fish failed to learn in the first block, it was categorized as a non-learner and was not considered further for the second training block. Overall, we behaviorally trained and recorded data from 26 larvae resulting in 39% learners, 46% non-learners and 15% intermediate learners, which were not included in further analyses.

#### Pre-processing of the neuronal signals

Each neuronal time series was de-trended individually to correct for the photo-bleaching, and then normalized as ΔF/F0 = (F-F0)/F0, where F is the fluorescence of the neuron at a given time point and F0 is the average fluorescence of the neuron across the entire recording. Noise was removed via total variation regularization by calculating the cumulative sum of the de-noised time derivatives (Chartrand, 2011). The de-noised time series X of each neuron were then normalized by taking the Z-score, given as (X – mean(X))/SD(X), for further analysis. This pre-processing procedure and subsequent analyses were performed using MATLAB (MathWorks).

#### Behavior Classification

Beside online tracking, offline behavioral classification was used to classify the tail movements as left turns, right turns, and struggles (or swimming), using a method similar to the one described by (Haesemeyer et al., 2018). Briefly, since larval zebrafish move in bouts rather than swim continuously, and since the bout duration lasts for about 250-400 ms in head-restrained fish (Severi et al., 2014), the movement bout was detected by scanning through the time series of the tracked tail angle in a sliding window of 50 frames (corresponding to ∼312 ms at a frame rate of 160 Hz). Within the time window of one bout, the time of movement initiation was set to the time point when the tail angle exceeded 5° for the first time. Then, the the bias of tail movements was calculated and the bout was categorized as a unilateral ‘turn’ or a bilateral ‘struggle’ with the threshold of bias at 1.05. The turn direction was determined from the sign of the tail angle, negative or positive, for left and right turns (labelled “TurnL” and “TurnR”), respectively.

#### Reaction Time

Reaction time (RT) was measured as the time between heat onset to the initiation of the first turn, regardless of correct or incorrect outcome, in a given trial. Because the temperature changes from the heat stimulus became relatively stable 5 s after heat onset, only trials with RT longer than 5 s were used, to minimize stimulus-related variation across trials.

#### Partial Correlation Coefficients

The partial correlation coefficient r of X and Y while controlling for the effect of Z is given as:

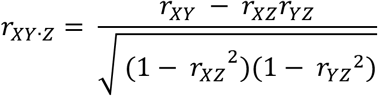

Here, *r_XY_* is the pair-wise correlation coefficient:

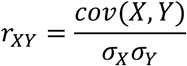

Partial correlation coefficients were computed in MATLAB (MathWorks), using the function *partialcorr()*.

#### t-SNE on the brain state

t-Distributed Stochastic Neighbor Embedding (t-SNE) (Maaten and Hinton, 2008) was applied to the sequence of brain states, where each “brain state” input vector was the activity state of all neurons at a given time point. 25,000 to 35,000 input vectors were mapped onto a 2D t-SNE map using value of 1000 for the “perplexity”, which is a critical hyper-parameter in t-SNE. Other values for perplexity, such as 500-2000, gave similar results. t-SNE maps from different animals were normalized to [0,1] in both the x- and y-dimension. Pair-wise distances of all pairs of brain states were measured by their Euclidian distance in the 2D t-SNE map.

#### t-SNE on the neuronal space

t-SNE was applied to the “neuronal space”, where each input vector was the normalized time series of one neuron in one recording, plus the four behavior- and stimulus regressors. 4000 to 6000 inputs were transformed into a 2D t-SNE map. Perplexity values ranging from 40 to 120 were tested and then chosen such that in the resulting map the regressors were mostly well separated. Hierarchical clustering was then performed on the chosen t-SNE map using the criterion of Ward’s minimum variance method (Ward, 2012). The final number of clusters was determined as the smallest number of clusters such that the top three principal components of each cluster explained more than 80% of the variance, or the first component more than 50%, using principal component analysis (PCA).

#### Identification of the neurons of cerebellum and ARTR for further analysis

Due to individual differences in neuronal activity, the neurons in the cerebellum and ARTR did not always end up in similar locations in the t-SNE neuronal spaces across animals. To ensure that the analysis was always performed on the medial portion of cerebellum and ARTR across animals (Figure 5-7), the neurons in these regions were selected manually based on anatomical landmarks (Dunn et al., 2016; Randlett et al., 2015; Takeuchi et al., 2015).

#### Population Dynamics using Demixed PCA (dPCA)

dPCA was performed on neuronal populations, following instructions described previously (Kobak et al., 2016). The goal of dPCA here was to project population dynamics into a small number of dimensions that capture more than 80% of the variance and to decode decision-dependent versus decision-independent components.

Briefly, for each neuron in a given recording, the time series was divided into correct and incorrect trials based on whether the first turn was in the correct direction. To be able to average trials of different reaction times (RTs) in dPCA, time series from different trials were first aligned relative to two time points, 5 s after heat onset and after movement initiation, respectively, using a time warping process, with RT 20 s (a.u.) after alignment (Figure S3). Note that only trials with RT ≥ 5 s were used for dPCA. The time window of each trial started 10 s before heat onset and ended 20 s after the first turn initiation. For each neuron in each trial, the average activity during the first 10 s was used as the baseline and subtracted from the time series. Because the movements were the only differing conditions across trials, two types of components were defined: decision-dependent components, from which the correct vs. incorrect trials could be decoded, and decision-dependent components, in which the population dynamics in correct and incorrect trials followed the same patterns.

To examine whether individual decision-dependent dPCs could decode correct versus incorrect trials with statistical significance, we used each dPC as a linear decoder to assess their classification performance (Kobak et al., 2016). Cross-validation (n = 100) was used to measure time-dependent classification accuracy of the dPCs. Then a shuffling test (n = 100), which shuffled the identity of correct vs. incorrect trials without changing the neuronal time series, was used to assess whether the classification accuracy was significantly above 99% of the shuffled runs in 10 consecutive time points. Time points of significant decoding were marked with thick horizontal black lines in Figure 4D, 5E, 5H. The dPCs which were able to decode correct vs. incorrect trials before movement initiation were defined as pre-motor decision-dependent components.

#### Analysis of Pre-motor decision-dependent activity

To compare the ability for predicting decision directions between learners and non-learners, warped time series with an RT of 20 s were used and the same procedure as above was performed.

To measure the time point of successfully predicted decision direction in learners, first warped time series were used to obtain the decoder of dPCA and the classification outcome from single-trials via cross-validation. Classification outcomes of single trials then went through an unwarping process by re-stretching back to the original RTs. Classification accuracy from unwarped single-trial classification outcomes was obtained by aligning to the movement initiation time. Error margins of classification accuracy were obtained from a shuffling test (n = 100) using the same procedure and were aligned to the movement initiation time after shuffling. Time for predicting decision directions was defined as the earliest time point before movement initiation when the classification accuracy was significantly above 99% of shuffled runs in 10 consecutive time points.

#### Analysis of Ramping activity

The first dPC of the cerebellum and ARTR showed a ramping temporal activity during the period from heat onset to movement initiation across animals. To measure the velocity of the ramping activity, i.e. the ramping rate, dPCA was first applied to neurons in the cerebellum and ARTR from both hemispheres to obtain the decoder and encoder, and then the decoder from dPCA (Figure S4B) was used to transform the neuronal activity (without time warping) into the low-dimensional map space, then a linear fit was applied onto the first dPC from heat onset to 1 s before movement initiation in each trial, thus obtaining the ramping rate at the single-trial level.

#### Analysis of ipsi-contra interaction

To examine whether there existed interaction and competition between hemispheres, dPCA was applied to neurons in the cerebellum and ARTR from both hemispheres to obtain the decoder and encoder, and then this decoder from dPCA was used to transform the neuronal activity (without time warping) into the low dimensional space. Since the response of each neuron can in principle be tuned to both features of the stimulus and of the motor output, a bilateral differential activity between the hemispheres during the pre-motor period may be masked by other stronger signals in the cerebellum and ARTR, such as the decision-independent dPC1. Thus, to increase the sensitivity of the analysis, neuronal activity containing only the pre-motor decision-dependent component was used. To do so, only the pre-motor decision-dependent component with the corresponding encoder was used to reconstruct the neuronal activity. With these two linear transformations, neuronal activity was obtained that was predictive for movement direction with temporal and spatial information. Then the neurons were pooled according to ipsilateral or contralateral location based on their anatomical locations in the fish brain, and their activity was averaged across pools. A time window from −4 to −2 s (note that this was the unwrapped time) before turn initiation was used, which was within the time window of 5 s RT as the threshold. The separation of 2 s from turn initiation was chosen to avoid any influence of post-turn activity from the de-noising process since de-noising by total variation regularization leads to a smoothing of the time series. The averaged activity of the ipsilateral and contralateral cerebellum and ARTR were then compared at the single-trial level.

### QUANTIFICATION AND STATISTICAL ANALYSIS

A total of 26 larvae were trained under the ROAST assay and 5 larvae went through spontaneous movement. LFM data from two-block learners, non-learners and fish exhibiting spontaneous movements - but not the data from intermediate learners - went through the 3D reconstruction. Subsequently, our signal extraction pipeline was applied to the reconstructed LFM images. We found imaging results and data quality to be reliably reproducible and consistent, both across imaging sessions with the same animal, and across animals. All statistical tests are described in the corresponding figure captions, and non-parametric statistical tests were used throughout this study as we did not assume normality of the data.

### DATA AND SOFTWARE AVAILABILITY

Our custom data-processing pipeline is available on GitHub. Data is available from the corresponding author upon reasonable request.

## SUPPLEMENTAL FIGURE TITLES AND LEGENDS

**Figure S1. Related to Figure 1.**
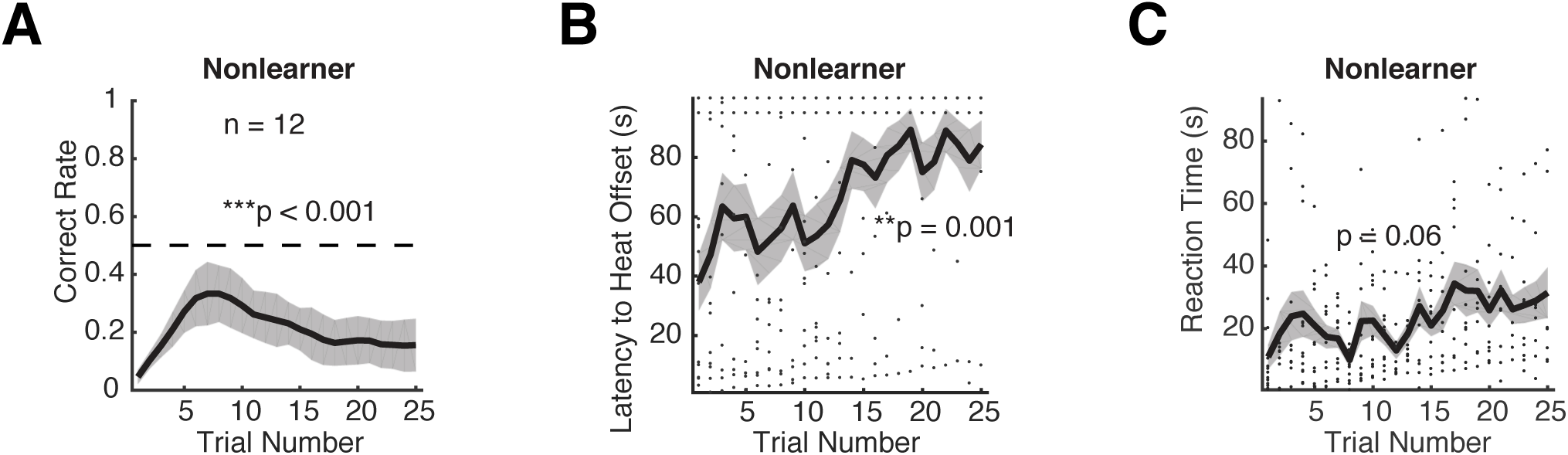
Performance of non-learners in the ROAST assay. (A) Performance of non-learners as a function of trial number. Moving average of correct rate over 3 trials (mean ± SEM). 7-8 dpf larvae, n = 12, siblings of learners. The correct rate for the last 10 trials is significantly below 0.5 (***p < 0.001, bootstrapping n = 5000), with a decreasing trend. (B) Increase of the heat duration (mean ± SEM) as a function of trials from non-learners. Two-tailed **p = 0.001, Kruskal-Wallis test. Same animals as in (C). Black dots are individual trials. (C) No significant change in the reaction time (mean ± SEM) of non-learners, as a function of trial number. Same animals as in (B). Two tailed p = 0.06, Kruskal-Wallis test. Black dots are individual trials.

**Figure S2. Related to Figure 2.**
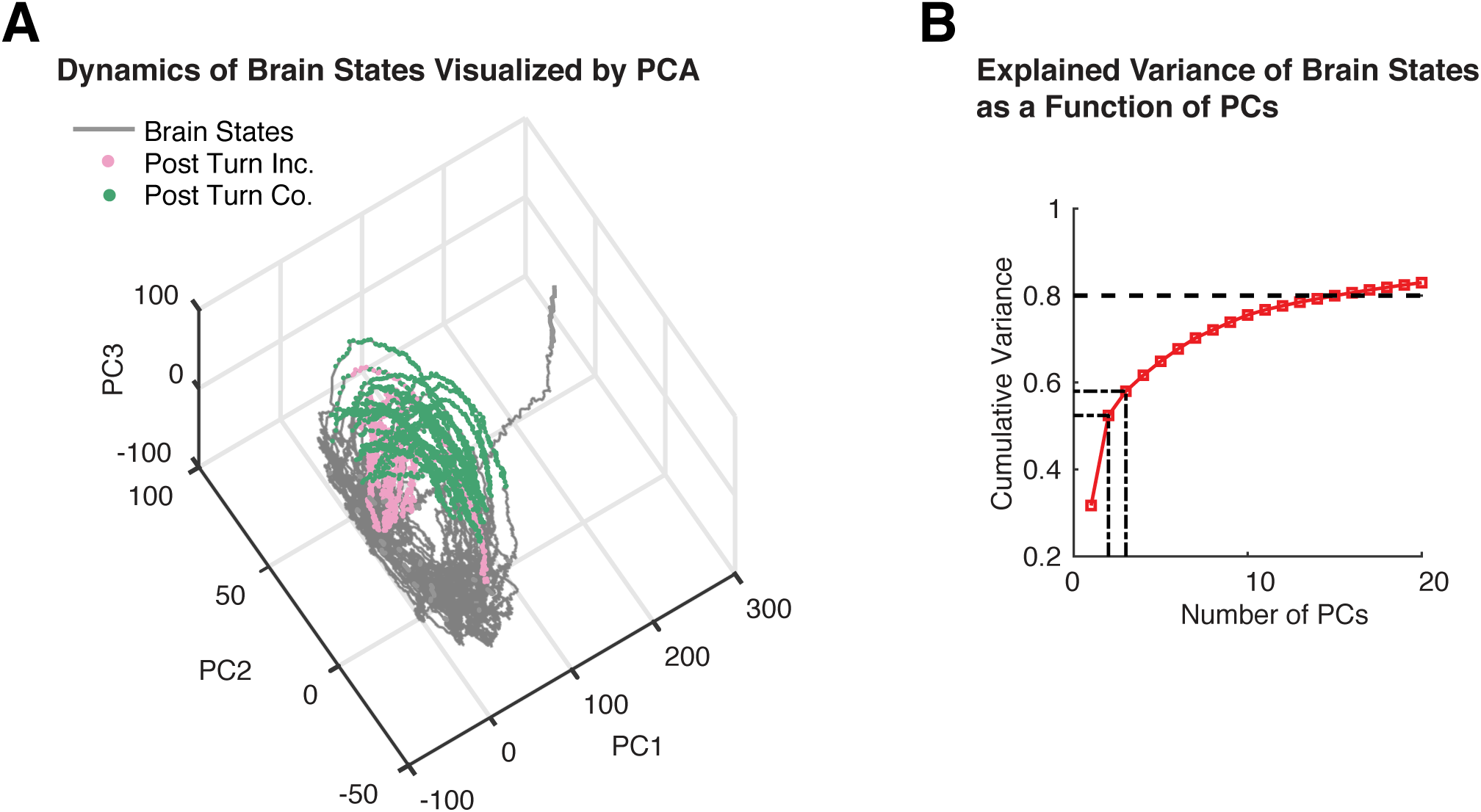
Ineffective separation of neural dynamics at different task epochs by principal component analysis (PCA). (A) Brain states represented as the top three principal components as a function of time. Same data as Figure 2. Viewing angle is chosen in attempt to distinguish post-correct (Co.) versus post-incorrect (Inc.) turns, which however is not possible. As the brain states are high-dimensional, only visualizing the top-ranking PCs is not informative, unless lower ranking PCs are further examined in separate plots. An alternative method for dimensionality reduction and visualization is required. (B) Cumulative explained variance of brain states as a function of number of principal components included (PCs). The brain states are high-dimensional, since explaining 80% of the total variance requires 15 dimensions.

**Figure S3. Related to Figure 3.**
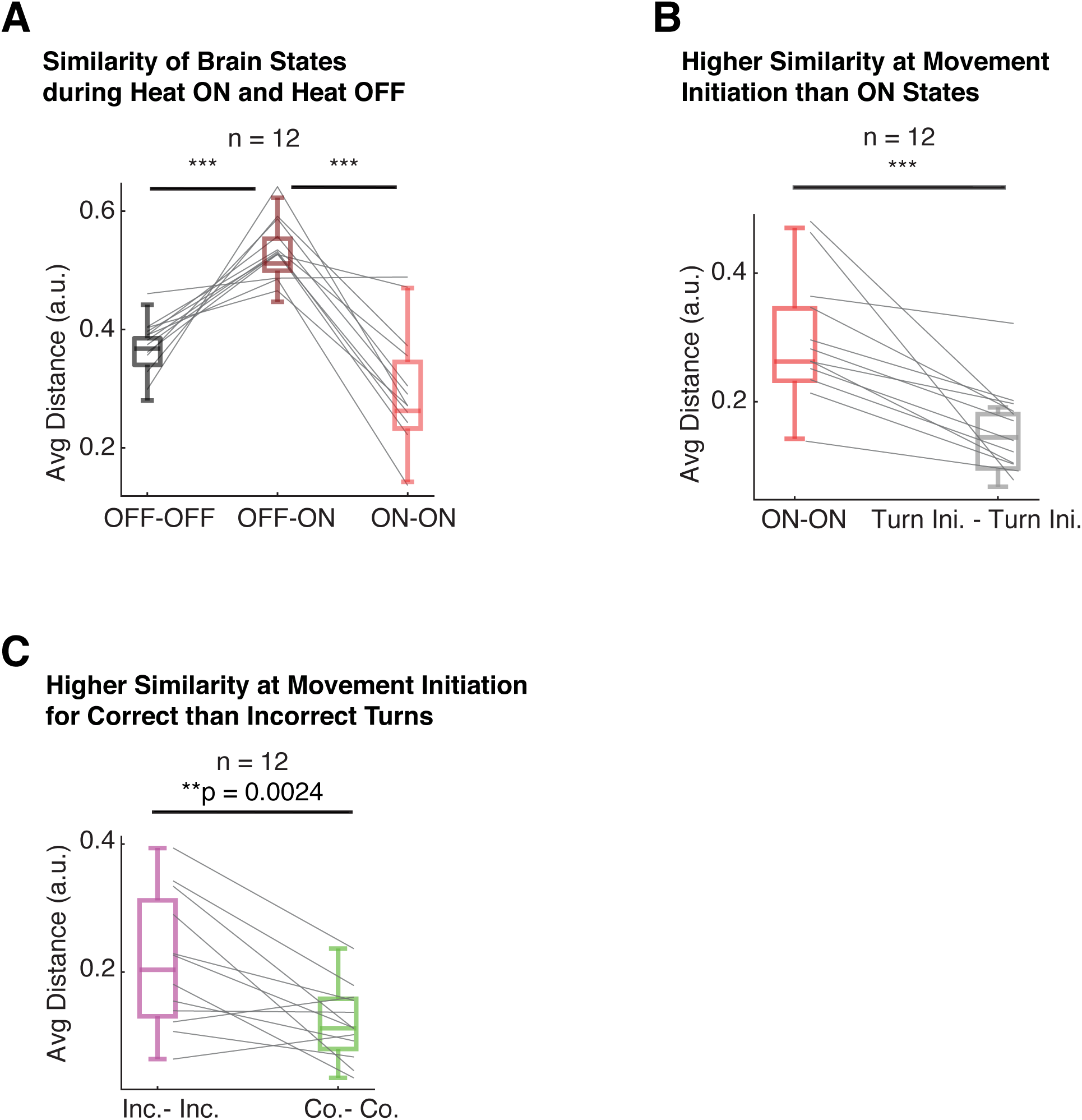
Dynamics of the brain states exhibiting recurrence to stimuli and bifurcation to behavioral outcomes. (A) Comparison of distance between brain states during Heat ON versus Heat OFF. Within-epoch brain states show high similarity than cross-epoch, consistent with Figure 3B. Distance between brain states is averaged over individual blocks and each pair of data is from same block. Data are from the 12 blocks of 6 learners, using Wilcoxon signed rank test, one-tailed, ***p < 0.001, same for other panels. (B) Comparison of distance between brain states during Heat ON versus Turn Initiation. Distance between brain states is averaged across individual blocks and each pair of data is from same block. Consistent with the representative data in Figure 3D, brain states at turn initiation are located closely in the “Heat ON” cluster. (C) Comparison of distance between brain states at correct versus incorrect turn initiation. Distance between brain states is averaged over individual blocks and each pair of data is from same block. Consistent with the representative data in Figure 3E, brain states are more similar at correct turn initiations than at incorrect ones.

**Figure S4. Related to Figure 4.**
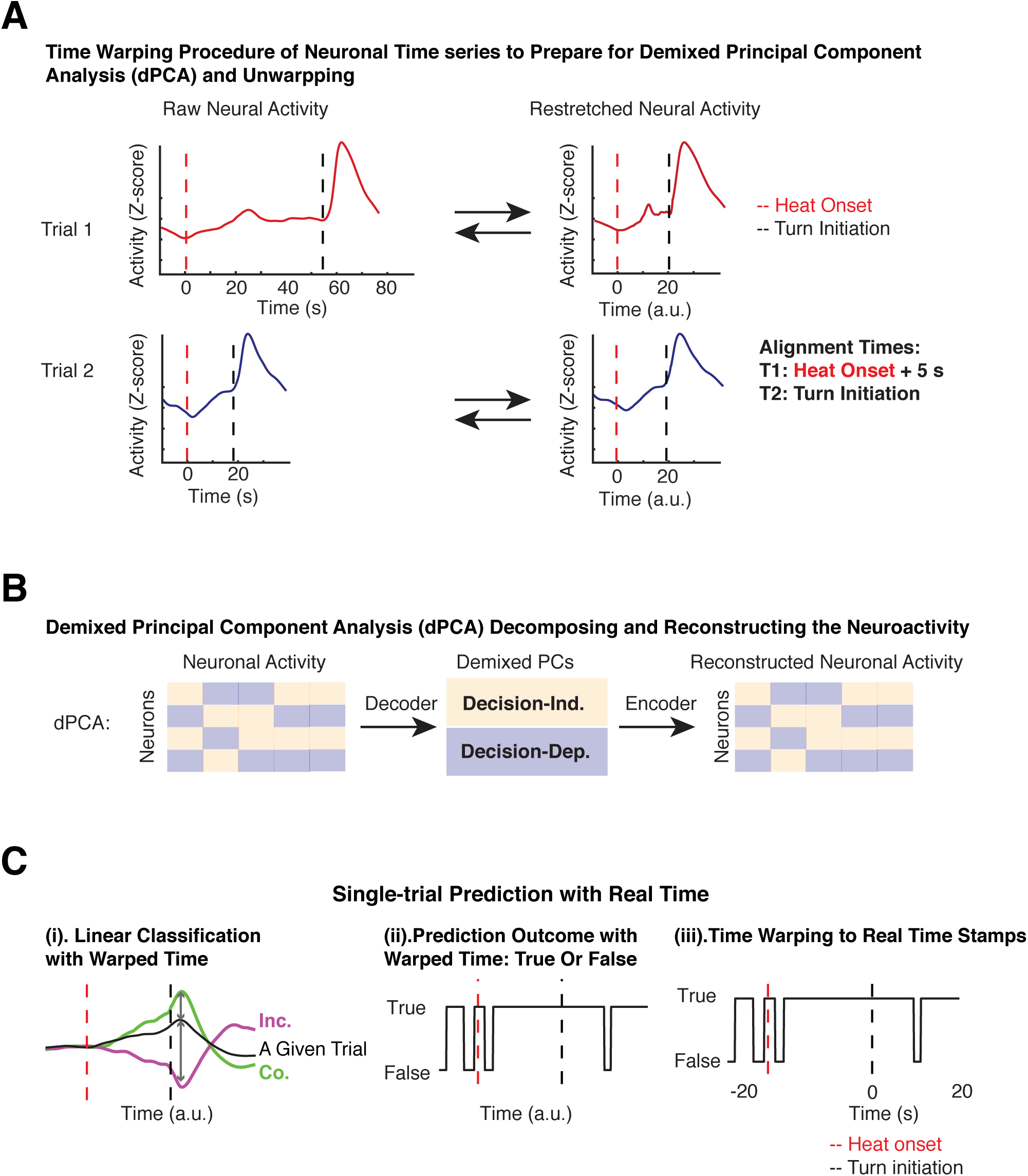
Illustration of the workflow of applying demixed PCA (dPCA). (A) Time warping procedure to align trials with different reaction times (RTs) for dPCA. To prepare the neuronal time series from different trials for dPCA, we define two alignment time points – t1, 5 seconds after heat onset, and t2 at turn initiation. These time points are chosen because the temperature becomes relatively stable 5 seconds after heat onset and to strictly avoid any contaminations from signals during post-turn period. For each trial, we find these two time points t1 and t2, and stretch or compress the neural activity between t1 and t2 to 15 seconds, thus making RT = 20 arbitrary units (a.u.), which is chosen based on the average RT. After this warping procedure, neural time series from different trials can be analyzed by dPCA to obtain the decoder and encoder. Furthermore, as the RT for each trial is known, this process is reversible, and demixed principal components (dPCs) can be stretched back to the original, i.e., unwarped time. For prediction of time, RT-related and ramping-related analyses (such as Figure 5H, 6 and 7), we use real time stamps instead of the warped ones. (B) Flowchart for decomposing neuronal population activity into task-related components, i.e. decision-dependent and decision-independent, by dPCA. This flowchart illustrates the process of compressing and decompressing the population activity through two linear transformations. The compressing step via the decoder provides the latent dPCs, the decomposed low-dimensional population dynamics. The decompressing step via the encoder transforms dPCs back to the space of neuronal time series. (C) Illustration of single-trial prediction for decision direction using real time by an unwarping procedure. (i). Classification of decision direction as a function of time using warped time with a linear classifier on the decision-dependent dPCs. Green and magenta curves: trial-averaged dPC from trials as the training data, correct and incorrect trials, respectively. Black curve: single-trial dPC from the test trial. Classification is by comparing distance between the test trial and correct versus incorrect trials at each time points. (ii). Classification outcome, true or false, at each time point of the test trial, in warped time. (iii) Real time prediction outcome, using a second time warping process which is reversed to panel (A).

**Figure S5. Related to Figure 4.**
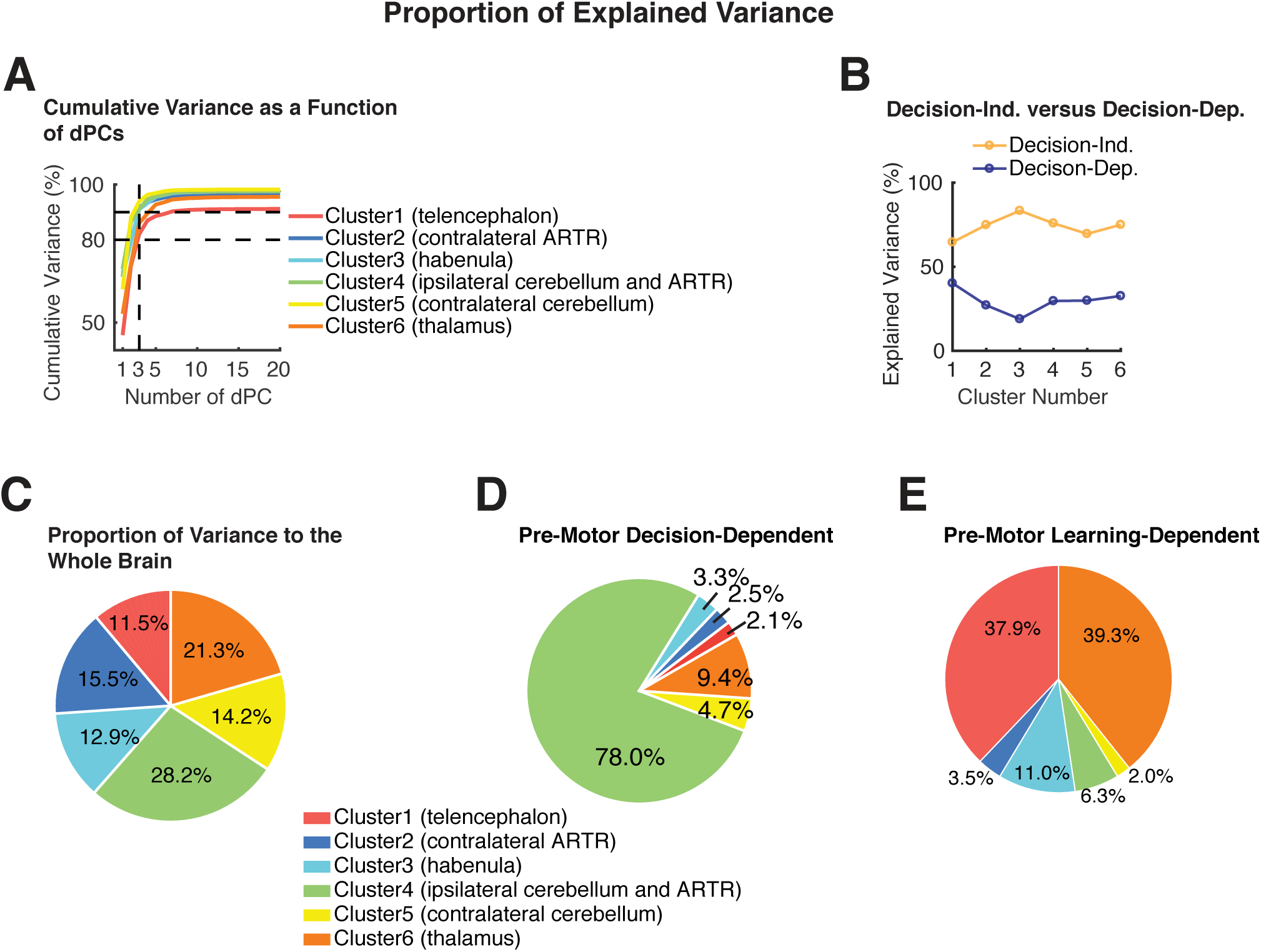
Proportion of explained variance of individual clusters and components across the whole brain activity. (A) Within each cluster, cumulative variance explained by dPCs. The top 20 dPCs are assessed here, which take up 91-98% of the total variance of a given cluster. The horizontal dashed line marks 90% and 80% variance. The top 3 dPCs, marked by the vertical dashed line, take up >80% of the variance of any given cluster. (B) Within each cluster, proportion of explained variance by decision-dependent versus decision-independent components. These two categories of components are not orthogonal to each other, i.e. partially correlated, so their sum exceeds 100%. This is because dPCA does not impose an orthogonality constraint when finding a decoder and encoder for different task-related parameters (Kobak et al., 2016). (C) Across the whole brain, proportion of explained variance by each cluster. (D) For pre-motor decision-dependent activity across the whole brain, proportion of variance from each cluster. Cluster 4, the ipsilateral cerebellum and ARTR, displays the largest portion (78%), even though its overall variance only takes up 28.2% of the entire brain. The second largest portion is with Cluster 6, the thalamus and sparse telencephalon neurons (9.4%). The third largest portion is with Cluster 5, the contralateral cerebellum (4.7%). Cluster 1 (telencephalon), Cluster 2 (the contralateral ARTR and lateral hindbrain) and Cluster 3 (the habenula and raphe) are also with pre-motor components, with proportion of each at 2-3%. Even though the pre-motor signal is available across the whole brain, the cerebellum and ARTR in total (Cluster 4 + Cluster 5) contain >80% of pre-motor components. (E) For pre-motor learning-dependent activity across the whole brain, proportion of variance from each cluster. Cluster1 (telencephalon) and Cluster 6 (thalamus) are with the largest portion (37.9% and 39.3% respectively), even though the overall variance of these clusters takes up only 11.5% and 21.3%, respectively, of the entire brain. The third largest portion is in Cluster 3, the habenula and raphe, 11.0%. For the cerebellum and ARTR, Cluster 4, Cluster 5 and Cluster 6, each is with <7% variance.

**Figure S6. Related to Figure 5.**
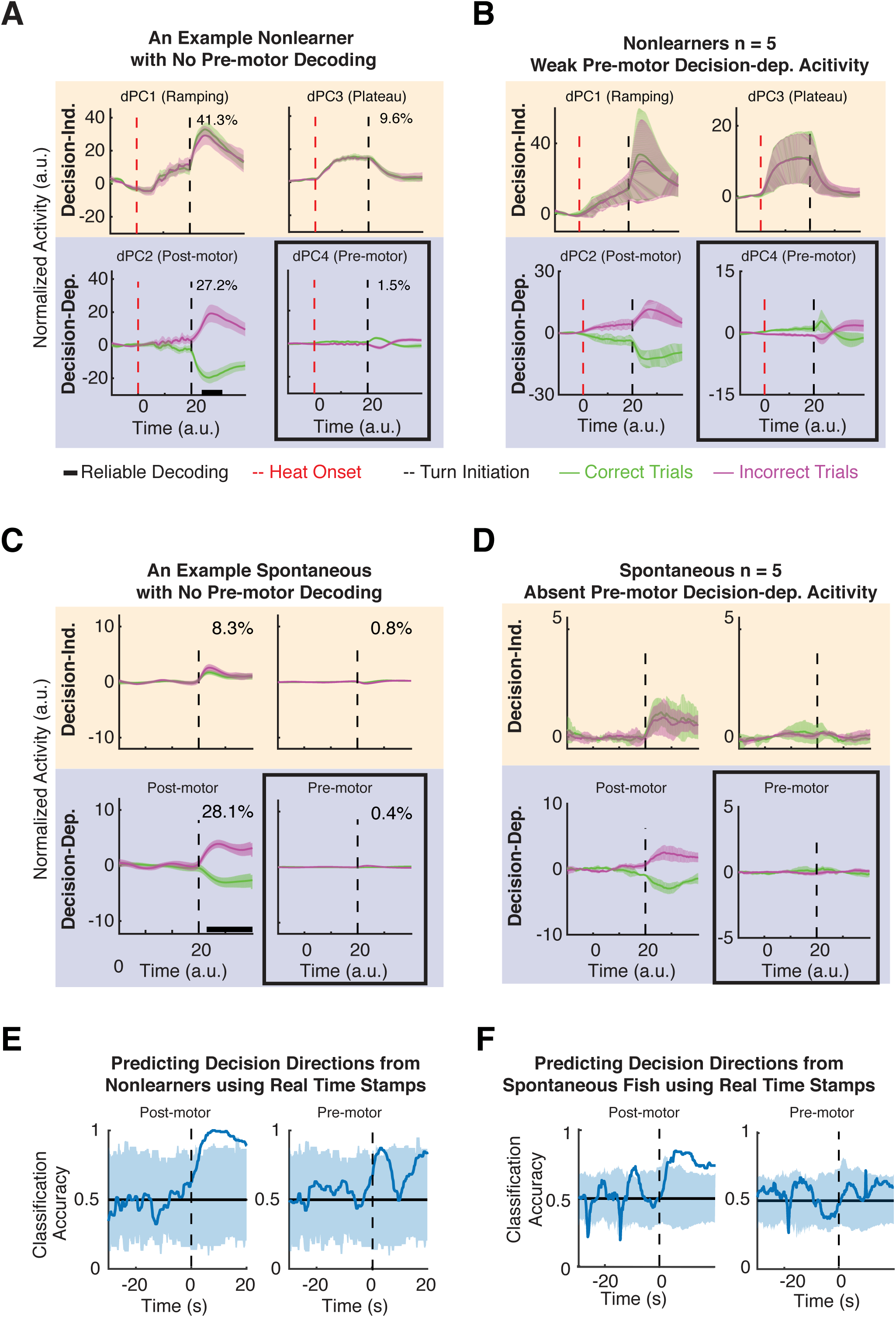
Decision-dependent and decision-independent dPCs in the cerebellum and ARTR from non-learners and fish under spontaneous movements. (A) Task-related dPCs from an example non-learner. Decision-independent dPC1 (heat-evoked ramping signal) and dPC3 (heat-and-motor-evoked signal) are similar to the example learner (Figure 5E); decision-dependent dPCs, the post-turn feedback signal as represented by dPC2, is similar to that of the learners (Figure 5E); dPC4, the pre-motor component, appears weaker than the learner (Figure 5E), and not predictive of the future turn direction. dPCs are displayed as mean ±95% CI across trials. (B) Averaged task-related dPCs across multiple non-learners. While the decision-independent dPC1 and dPC3 are similar to learners (Figure 5F), decision-dependent dPCs exhibit different patterns than learners. dPC2 displays post-motor difference between correct versus incorrect trial; dPC4, which is the pre-motor component in learners, displays weak activity with small margins between correct versus incorrect trials. (C) Task-related dPCs from an example fish performing spontaneous movements. No stimulus is delivered in this task. Decision-independent dPC1 and dPC3 are almost absent, without pre-motor ramping or plateau, while a weak post-motor separation by the dPC2 is present. Decision-dependent dPC4 is absent. (D) Averaged task-related dPCs from fish performing spontaneous movements. The decision-independent dPC1 and dPC3 are much weaker, and without pre-motor ramping or plateau. Decision-dependent dPC2 displays post-motor separation, similar to learners and non-learners, while dPC4 shows no pre-motor separation. (E) Time for predicting decision directions from non-learners using real time. No predictive signal is present. Similarly, in the other four non-learners that were analyzed, no predictive signals are present either. (F) Time for predicting decision directions from fish undergoing spontaneous movements, using real time. This fish does not show predictive decision-dependent dPCs. Similarly, in the other 4 fish that were analyzed, no predictive signals are present.

**Figure S7. Related to Figure 6.**
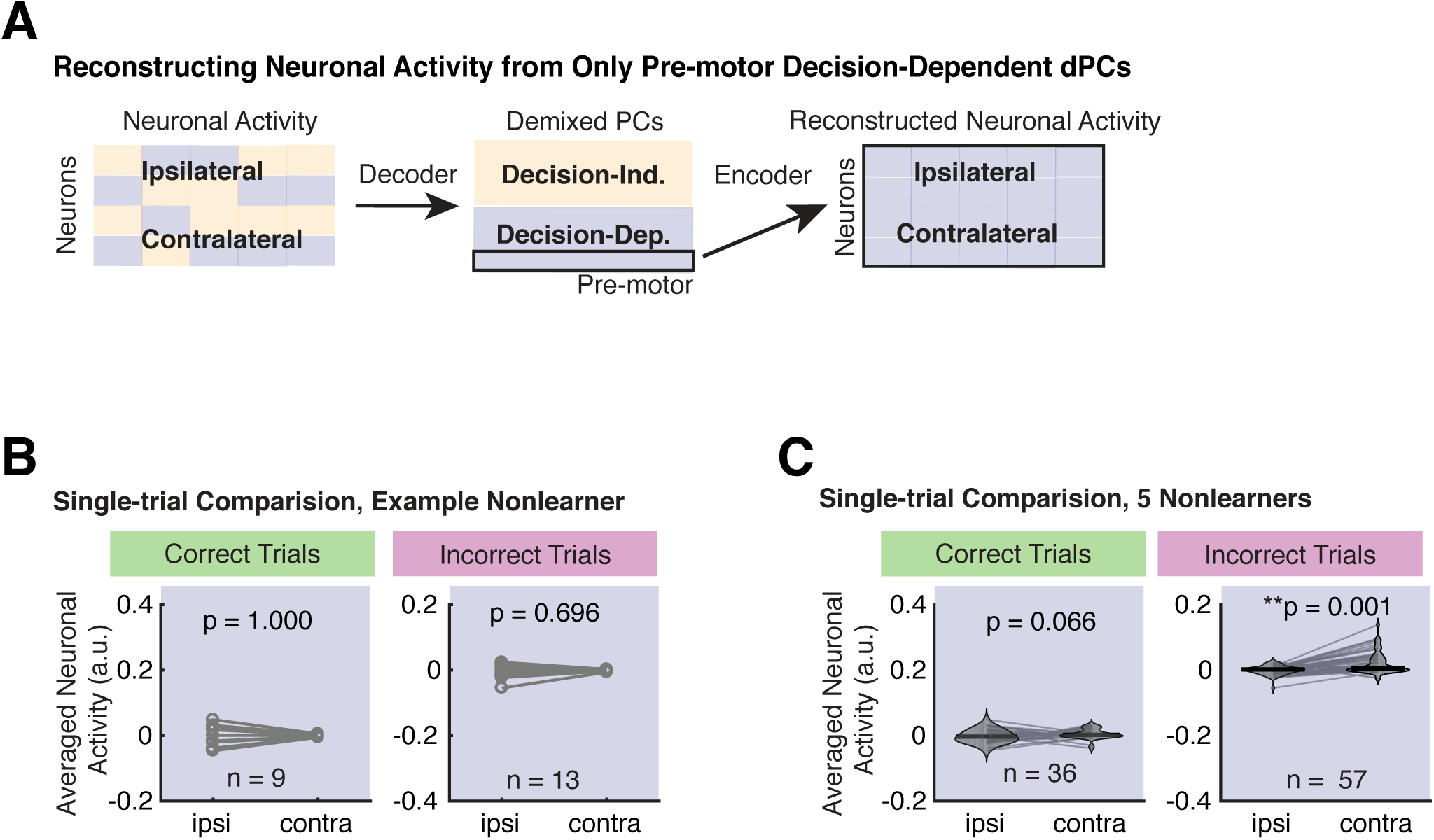
Weak and even absent ipsilateral-contralateral difference in the pre-motor neuroactivity of the cerebellum and ARTR in non-learners. (A) Illustration of the workflow for reconstructing neuronal activity from pre-motor decision-dependent dPCs only. dPC4 is used for reconstruction based on the results shown in Figure 5F. (B) Comparison of reconstructed neuronal activity during pre-motor periods between ipsilateral and contralateral sides on a trial-by-trial basis, from an example non-learner. Pre-motor activity is averaged from 4 to 2 s before turn initiation. Unlike the example learner, the neuronal activity on the ipsilateral side of the non-learner is not significantly different from the contralateral side in both correct and incorrect trials. Wilcoxon signed rank test, two-tailed. (C) Comparison of the ipsilateral-contralateral difference at the single-trial level from trials pooled across multiple non-learners. The ipsilateral pre-motor activity is not significantly different from the contralateral side during correct trials. Wilcoxon signed rank test, two-tailed.

**Figure S8. Related to Figure 7.**
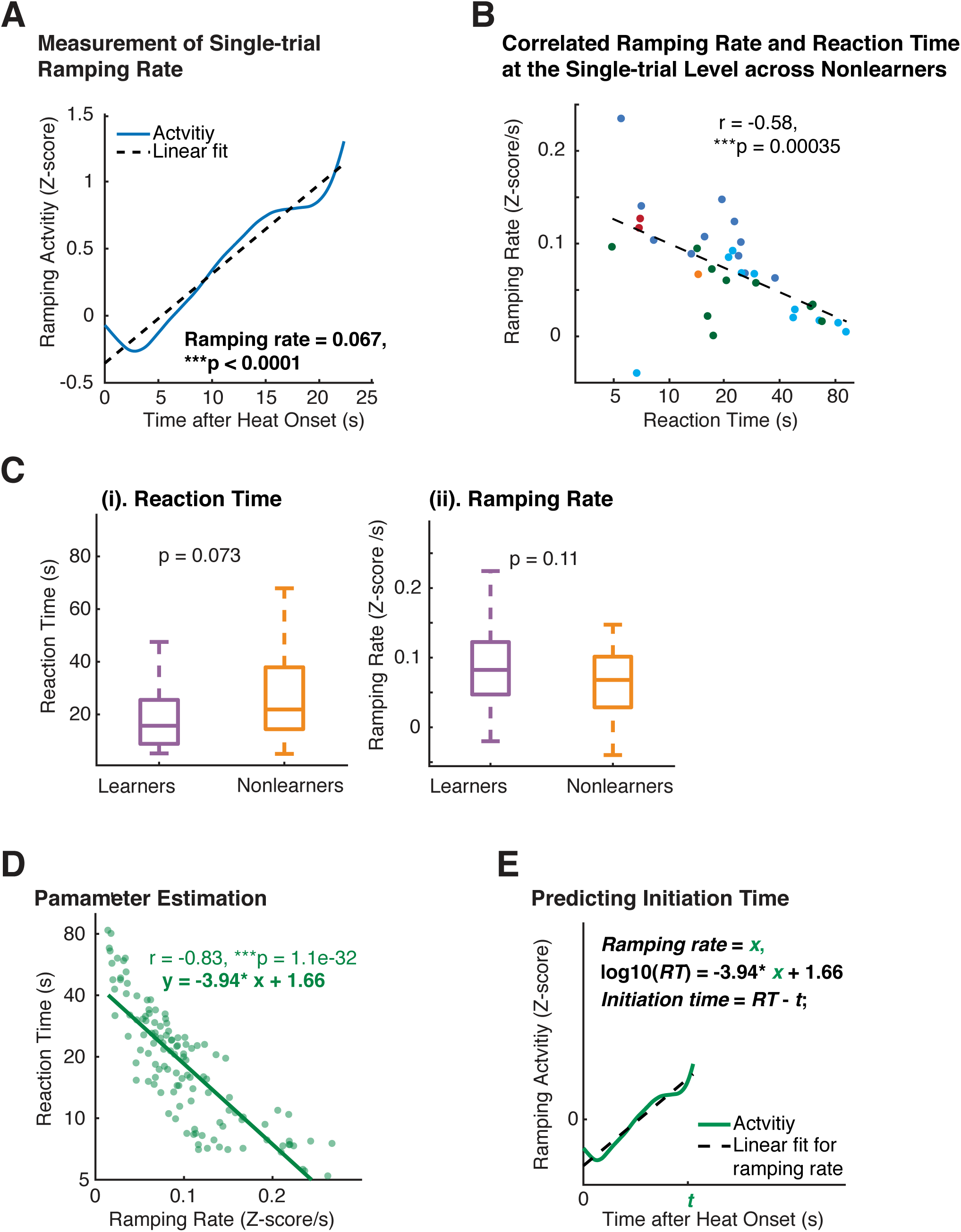
Single-trial ramping rate and comparison between learners and non-learners. (A) An example of how ramping rate of a given trial was evaluated. A linear fit is applied to the ramping activity from heat onset to 1 s before turn, using real time. The slope parameter is used as the ramping rate. (B) Inverse correlation between ramping rate of population activity from the cerebellum and ARTR and reaction time at the single-trial level in non-learners. Data is pooled across the correct trials of multiple non-learners. Each dot represents a single trial; each color indicates one fish. Dashed lines indicate linear regression. 5 training blocks from 5 non-learners are shown here. The correlation coefficient between ramping rates and reaction times from non-learners is similar to the one from learners in Figure 7D. Even though these animals fail to learn, the inverse relationship between reaction times and ramping activity is still highly significant. (C) No statistical difference in the ramping rates or reaction times between learners and non-learners. Wilcoxon rank sum test, two-tailed. Note that incorrect trials and trials with reaction time shorter than 5 s are not included in this analysis. (D) Estimating parameters for the linear equation between ramping rate and the logarithm of the reaction time. Note that trials were defined as outliers when their ramping rate × log10(RT) was <0.5. Outliers were not included in parameter estimation. Here we obtain the linear fit of RT as a function of ramping rate at the single-trial level. Each dot represents a single trial. Line indicates linear regression. (E) Illustration of the prediction of movement initiation time at a given time after heat onset in a given trial.

## SUPPLEMENTAL ITEM TITLES AND LEGENDS

**Movie S1. Projected volumetric reconstruction of whole-brain LFM calcium imaging data. Related to Figure 2.**

Maximum intensity projections along three orthogonal directions of reconstructed LFM recording of GECI activity in an example zebrafish with pan-neuronal expression of GCaMP6s. Scale bar, 50 µm. Recording frame rate: 10 fps. Playback: 30× real time.

**Movie S2. Single-trial bifurcation process during the pre-motor period. Related to Figure 3I.**

Animation of the evolution of individual correct and incorrect trials during the pre-motor period (i.e., from heat onset to movement initiations), expressed as correlation with the averaged, final brain states at movement initiation. Black diamonds indicate heat onset, green and magenta dots indicate movement initiations for correct and incorrect turns, respectively. Color scale encodes temporal progress from heat onset to movement initiation. Note that these individual trials are with different reaction times. Playback at 3× real time.

**Movie S3. Predicting turn direction and turn time from neural activity at the single-trial level. Related to Figure 5-8.**

Movie of calcium imaging data during a ROAST trial, together with evolution of prediction for both, turn direction and turn time. Prediction is obtained from the neuronal activity at the single trial level. The example is a correct trial, with the training towards the right direction. Video starts 5 s before heat onset and ends 5 s after movement initiation. The prediction is evaluated starting at heat onset and updated until movement initiation.

Top panels: Maximum intensity projections of neuron centroids, superimposed on an anatomical reference, *Tg(elavl3:H2B-RFP)* from the Z-Brain Atlas (Randlett et al., 2015). Filled red square indicates Heat ON, hollow red square indicates Heat OFF. Scale bar, 50 µm.

Middle panel: Outcome of a binary prediction for turn direction. Hollow triangles indicate the possible outcomes (left versus right), and filled triangles indicate predicted outcome, color coded green for the right (correct) side and magenta for the left (incorrect) side.

Bottom left panel: Evolution of error in predicted turn time, i.e. the difference between the prediction and the actual turn time. Red dashed line indicates heat onset; white dashed line indicates turn initiation.

Bottom right panel: Plot of tracked tail segments (n = 9 segments). Each rendered frame is the maximum-projection of 16 consecutive recorded frames, to match behavioral recording frame rate (160 fps) to brain imaging frame rate (10 Hz). Playback is in real time (10 fps).

